# HiFIseek: gene-specific enrichment of high-impact mutations in associated genomic regions

**DOI:** 10.1101/2025.06.12.659305

**Authors:** Rafael Riudavets Puig, Ina Skaara Brorson, Denise O’Mahony, Vessela N. Kristensen, Anthony Mathelier

## Abstract

Transcription is regulated through the sequence-specific binding of transcription factors (TFs) to cis-regulatory regions (CRRs). Although scattered along the genome, multiple CRRs are brought in close spatial proximity to the genes they regulate through the formation of DNA loops. The sequence component of transcriptional regulation suggests that DNA variants in CRRs could disrupt TF-DNA interactions and gene regulatory networks through a cascading effect. To date, only a few cases of recurrent cis-regulatory variants have been described. As an alternative to variant recurrence, some methods utilize genomic annotations that indicate the functional impact (FI) of a variant and the CRRs of each gene to detect the enrichment of high-impact cis-regulatory variants (CRVs). However, the agreement across the different associations and FI scoring methods remains unexplored. This work demonstrates that gene-CRR and FI scoring methods exhibit little consensus, highlighting the impact of the choice of a specific CRR-score combination in a given analysis. In addition, we demonstrate that cancer genes exhibit a higher frequency of FI variants compared to non-cancer genes. Based on this, we developed HiFIseek, a Snakemake pipeline exploring the enrichment of FI variants in associated genomic regions using all region-score combinations and two enrichment detection methods. We apply HiFIseek to detect genes exhibiting high FI CRVs in eight cancer cohorts and a set of breast cancer risk-associated single-nucleotide polymorphisms (SNPs). In the cancer cohorts, we observe a small degree of agreement across CRR-score combinations. Although HiFIseek detected known cancer-related genes with FI CRVs, most of them did not show significant changes in their expression. In the SNPs cohort, HiFIseek found small consensus across CRR-score combinations. However, one of the combinations returned 18 known cancer-related genes, including *BRCA1*, showing an enrichment of high-impact CRVs.

## Introduction

Transcriptional regulation is a highly complex mechanism involving several biological processes. One of its most essential elements consists of the interplay between transcription factors (TFs) and cis-regulatory regions (CRRs) (1). TFs are DNA-binding proteins with a selective affinity for short DNA sequences known as their TF binding sites (TFBSs). CRRs are genomic regions that play a crucial role in transcriptional regulation. Depending on their effect and location relative to the transcription start sites of their transcriptional targets, CRRs are classified into promoters, enhancers, and silencers. While promoters are located directly upstream of their transcriptional targets, enhancers and silencers can be found hundreds of kilobases away and in any orientation. Enhancers and silencers are involved in transcriptional activation and repression, respectively. The interplay between TFs and CRRs is achieved through the presence of TFBSs in CRRs, enabling their context-specific interaction. Because this interaction is sequence-dependent, some DNA mutations may alter transcriptional regulation.

Although many cancer driver mutations are mapped to protein-coding regions, the vast majority of somatic mutations fall within the non-protein-coding portion of the genome (2), where most of them are thought to be passenger events. Similarly, most disease-associated single-nucleotide polymorphisms (SNPs) are found in non-protein-coding regions (3). These observations suggest there is potential for driver events outside the protein-coding genome. However, understanding which of these variants are drivers and their underlying mechanisms is an ongoing challenge. The difficulty in detecting these events can be attributed to many reasons. For example, the interpretation of non-protein-coding variants is significantly more challenging than it is for their protein-coding counterpart. This is mainly due to the complexity of gene regulation, where the combinatorics of TF-CRR-gene interactions suggest that there may be many possible ways to disrupt a gene’s cis-regulatory space. Additionally, the impact of the same variant can be larger or lower depending on factors such as cell type and state (4).

Variant recurrence is commonly used to identify variants that show signs of positive selection in disease. Currently, only a handful of recurring events have been attributed to cis-regulatory variants (CRVs) in the context of cancer (5). The most well-known example involves the recurring mutations of the TERT promoter, which introduce new binding sites for TFs of the ETS family and lead to the reactivation of the TERT gene’s transcription (6, 7). Besides recurrence, other studies explored alternative ways to detect candidate non-protein-coding driver events. In (8), the authors studied the effect of CRVs on TF binding, showing some cases where the detected genes showed a significant change in gene expression. The study in (9) showed that CRRs bound by known prostate-cancer-related TFs such as AR, FOXA1, HOXB13, and SOX9 were enriched for CRVs and how these non-protein-coding mutations lead to the overexpression of MYC. Outside the context of cancer, the authors in (10) used promoter capture HiC (pc-HiC) in human primary hematopoietic cells to study which genomic regions are near gene promoters, finding an enrichment for expression quantitative trait loci (eQTLs) in pc-HiC derived CRRs. The work in (11) focused on obesity, showing how SNP rs1421085 disrupts the binding of ARID5B and results in a derepression of IRX3 and IRX5 during adipocyte differentiation.

Over the years, alternatives to mutation recurrence have been proposed to search for candidate driver events. One of these involves the search for the enrichment of functional mutations in sets of associated genomic regions. An example of this is OncodriveFML (12), which studies whether a set of genomic regions shows a higher-than-expected average impact score by comparing the observed average to a permutation-based expectation. However, averaging impact scores may mask the relevance of high-impact variants that are considered together with many low-impact ones. Additionally, permutation-based methods can be computationally intensive. Regardless of the methodology used to compute the enrichment, two key elements in this type of approach are the annotations grouping genomic regions and scoring the functional impact (FI) of mutations. For example, exploring gene CRRs for high-impact mutations requires knowing all relevant CRRs, their transcriptional targets, and the effect of mutations within them.

Dissecting which CRRs are relevant for the transcriptional regulation of each gene remains a challenging task. Although a broad range of methods attempts to tackle this problem, gold-standard methods and resources are still lacking in the field. The Genehancer (13) resource consists of a collection of known and predicted gene-CRR associations. Other methods use specific types of data to produce gene-CRR links. For example, epiregio gene-CRR associations are inferred by STITCHIT (14), which predicts gene expression using DNase I hypersensitivity sites (DHS). Another method is the activity-by-contact (ABC) model (15), which uses DHS, H3K27Ac, and chromatin conformation data to determine each gene’s CRRs while accounting for the activity level of each CRR. Other experimental methods such as pcHi-C (16) are specifically designed to obtain the genomic regions in close spatial proximity to a selected set of gene promoters.

Similarly to the case of gene-CRR associations, a gold-standard methodology scoring the functional impact of non-protein-coding variants is still lacking. The last years have seen the development of many different approaches assessing the functional impact of variants. Some approaches utilize available annotations to predict the effect of variants in a supervised manner. For example, the CADD (17) and DANN methods classify a variant as deleterious or neutral based on 60 different annotations using logistic regression and a deep learning framework, respectively. Similarly, the LINSIGHT (19) method combines a generalized linear model with a probabilistic graphical model to leverage molecular evolution and genomic data. While existing annotations enable the leveraging of current knowledge to infer variant impact, the FI of poorly annotated variants may be underappreciated. Likewise, the FI of non-protein-coding variants may be underappreciated if they are considered in conjunction with protein-coding ones. Other methods, such as EIGEN and EIGEN-PC (20), employ an unsupervised approach that combines different annotations. Common to these methods is their aim to provide one score for each possible variant in the genome, regardless of the cellular context. However, the same variant may have different impacts depending on the affected cell type or condition. This limitation is addressed in the Sei framework (21), which utilizes a deep learning model to predict TF binding and histone modifications from DNA sequences across multiple cellular contexts. The predicted chromatin profiles are then classified into sequence classes, some of which have been shown to represent specific cell types. Therefore, running an in silico mutagenesis approach while focusing on particular sequence classes can better measure the impact of variants in a cell type of interest.

Studies exploring variants in the cis-regulatory space of a gene typically rely on one chosen combination of association and variant scoring approaches. However, the impact of a chosen combination on the analysis results remains largely unexplored. Given the lack of gold-standard datasets and the context-specificity of transcriptional regulation, exploring all CRR-score combinations can be used to identify instances where multiple combinations agree. Cases showing an enrichment of high-impact CRVs across several CRR-score combinations can then be prioritized for further analysis.

In this work, we present evidence that gene-CRR association and FI scoring methods show a small level of consensus, and that cancer-related genes contain higher FI variants in their CRRs compared to non-cancer-related genes. Based on this observation, we further searched for the enrichment of high FI variants in each gene’s cis-regulatory space. To achieve this, we developed HiFIseek, a Snakemake (22) pipeline that explores any number of cohorts with all combinations of provided association and FI scoring methods. For each combination, HiFIseek computes the enrichment of high-impact variants using two different methods: (1) OncodriveFML, and (2) a new analytical approach to the OncodriveFML methodology. We apply this workflow to study each gene’s CRRs in seven cancer cohorts from The Cancer Genome Atlas (TCGA) (23), a breast cancer cohort from the International Cancer Genome Consortium (ICGC) (24), and a set of SNPs associated with increased risk of developing breast cancer from the Norwegian Breast Cancer Study (NBCS). We employed four distinct gene-CRR association methods and eight variant scoring methods. In the cancer cohorts, results varied across CRR-score combinations, with a total of 210 distinct genes across all cohorts showing an enrichment of high-impact CRVs in at least 5 different CRR-score combinations. Conversely, the application of HiFIseek to the set of breast cancer-related SNPs within gene CRRs resulted in well-known breast cancer-related genes, such as *BRCA1*.

## Results

### Gene-CRR association methods show small overlap

We conducted a series of exploratory analyses to gain a deeper understanding of the differences between gene-CRR association methods. We started by exploring the distributions of the sizes of each gene’s associated CRRs. In most association methods, region sizes showed an average size of ~1000 bp except Genehancer regions, which were found to be 1-2 orders of magnitude larger (Supplementary Figure 1). We then explored the agreement across association methods. For each gene, we calculated the proportion of overlap between its CRRs using different association methods. In many cases, we observed a small consensus across gene-CRR associations. Genehancer regions showed the largest overlap with all other region types. This may be due to two reasons: (1) these regions were part of the datasets used to compile Gene-hancer regions, and (2) the larger size of Genehancer regions. Among all other association types, ABC-derived gene CRRs showed the strongest overlap with all other associations except epiregio (Figure 1). We observed a strong agreement between ABC regions derived from different cell types (0.11 - 0.37 median overlap proportion). ABC regions also showed some overlap with intronic and promoter regions. Finally, epiregio regions did not overlap with any of the other gene-CRR association methods.

**Fig. 1.**
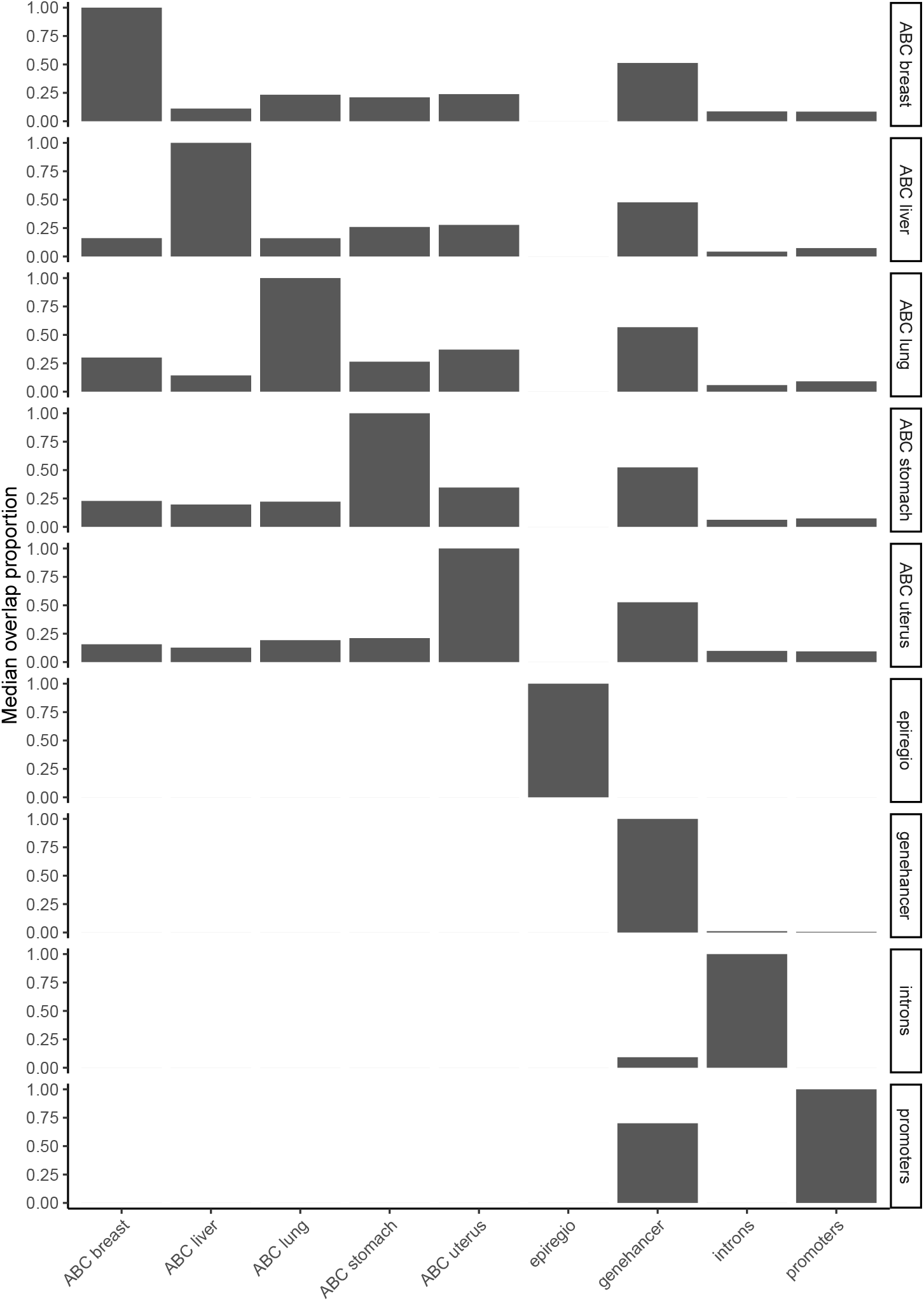
Gene-CRR association methods show small consensus. Median overlap proportion (y-axis) between gene-CRR association methods. Facet labels indicate the reference association method while the x-axis indicates the association method compared against. The median overlap proportion indicates the fraction of the reference association showing an overlap with the method being compared against.

In summary, these results show the lack of agreement across gene-CRR association methods, underscoring the significant impact that the method of choice can have when studying gene-CRR associations.

### Functional impact scores show little consensus

We next compared the distributions of variant impact scores across scoring methods. We observed that scoring approaches show substantial differences not only in the range of score values but also in their distribution (Supplementary Figure 2). For example, CADD and DANN scores utilize different models to analyze the same data and infer the impact of variants. However, their score distributions are vastly different.

We explored the correlation between methods when comparing the impact scores for the same set of variants. We randomly selected 10,000 distinct non-protein-coding mutations from all TCGA cohorts and compared their impact scores across all methods. Similar to the case of gene-CRR associations, scoring approaches showed little consensus on the impact of the selected mutations (Figure 2).

**Fig. 2.**
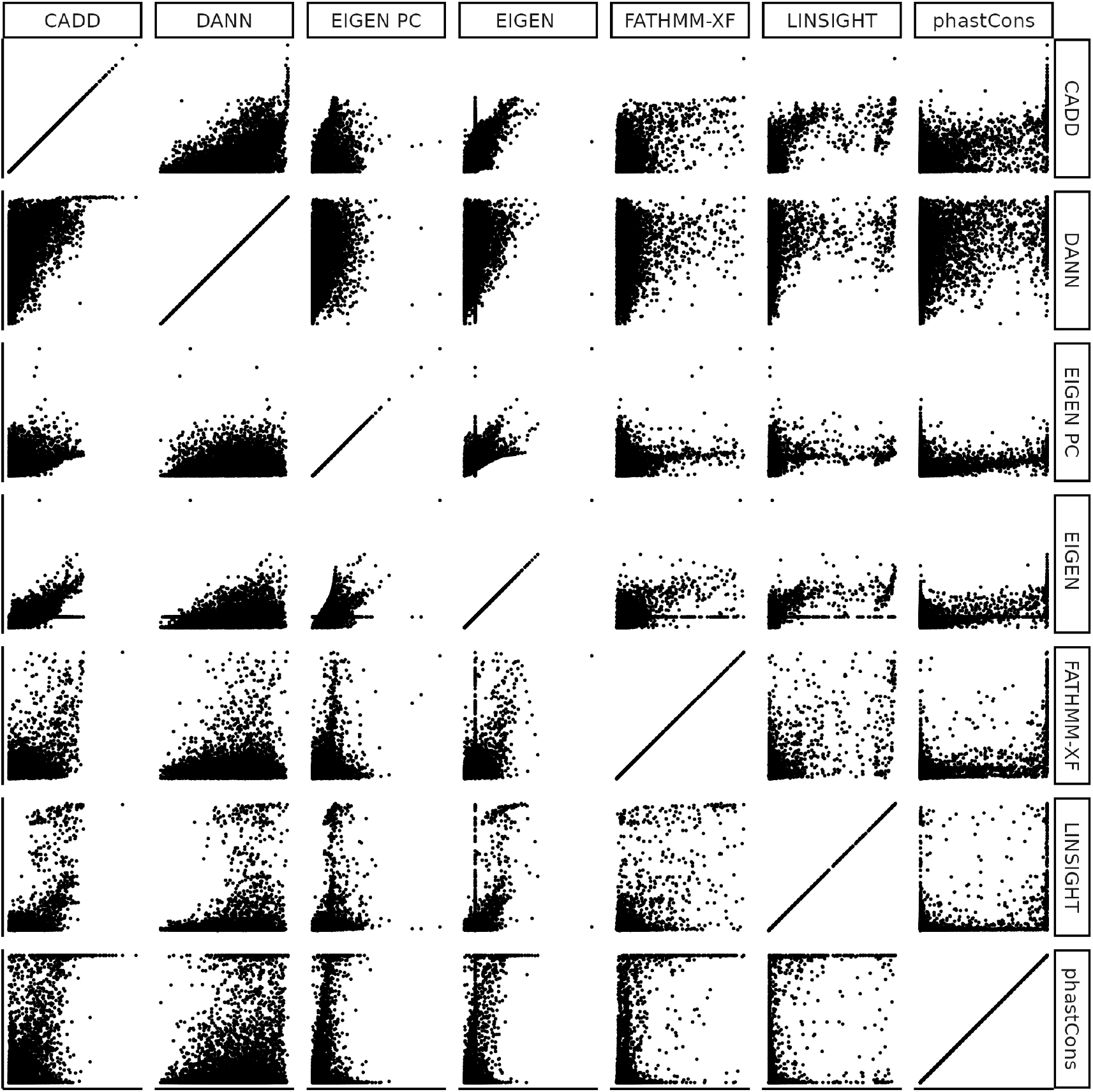
Variant impact scores show little agreement. Scatter plots comparing the scores for the same 10,000 randomly picked variants.

These results highlight the disagreement among FI scoring approaches when examining the potential impact of variants. Such differences are likely to affect the results of methodologies using these scores to detect enrichment of FI variants in a set of associated genomic regions.

### Cancer gene CRRs show significantly higher functional impact scores

We studied different variant patterns in data from seven TCGA cancer cohorts, the ICGC breast cancer cohort, and a set of SNPs associated with an increased risk of developing breast cancer from NBCS (Supplementary Figure 3). The studied TCGA cancer cohorts include: BRCA-US (breast invasive carcinoma), HNSC-US (head and neck squamous cell carcinoma), LIHC-US (liver hepatocellular carcinoma), LUAD-US (lung adenocarcinoma), LUSC-US (lung squamous cell carcinoma), STAD-US (stomach adenocarcinoma), and UCEC-US (uterine corpus endometrial carcinoma). We explored, for each cohort, the number of distinct mutations located in protein-coding or non-protein-coding regions. We utilized RefSeq annotations to investigate the number of distinct mutations located in gene promoters, 5’ UTRs, coding exons, introns, and 3’ UTRs. As expected, the majority of mutations occurred within the non-protein-coding genome, with intronic regions showing similar numbers of mutations to other non-protein-coding regions (Supplementary Figure 4).

We next explored the following properties for cancer versus non-cancer genes: (1) the distribution of gene region mutation rates stratified by association method; and (2) the distribution of FI scores for the observed mutations stratified by association and FI scoring methods.

We studied the distributions of mutation rates in the CRRs, promoters, 5’ UTRs, coding exons, introns, and 3’ UTRs of cancer and non-cancer genes. We used the following annotations to define gene-CRR associations: epiregio, Genehancer, and ABC when data for a similar cell type were available. Although cancer genes did not show significantly higher mutation rates in many of the studied regions, we observed significantly higher mutation rates in 5’ UTRs in all cancer cohorts, epiregio CRRs in the ICGC cohort, and ABC-derived CRRs in the TCGA LIHC cohort (Supplementary Figures 5-12).

We compared the FI scores for each cohort, distinguishing between cancer and non-cancer genes, across various association and scoring methods. We used the following FI scoring methods: CADD, DANN, EIGEN, EIGEN PC, FATHMM-XF, LINSIGHT, and phastCons across 100 vertebrate species. This analysis resulted in many region-score combinations where cancer genes showed significantly higher impact CRVs than non-cancer genes (Figure 3, Supplementary Figures 13-20). Promoter regions showed the strongest agreement across scores, where cancer genes showed significantly higher FI scores than non-cancer genes in virtually all scores and cohorts. Other region types, including ABC and Genehancer regions, showed more variation across scores and cohorts. Finally, cancer genes did not show higher FI CRVs when using epiregio CRRs in most cohort-score combinations except for the HNSC cohort. Based on these results, we decided to search for gene CRRs that enrich for high FI CRVs, using all combinations of CRR-score combinations, while focusing on those combinations where cancer genes show higher FI scores.

**Fig. 3.**
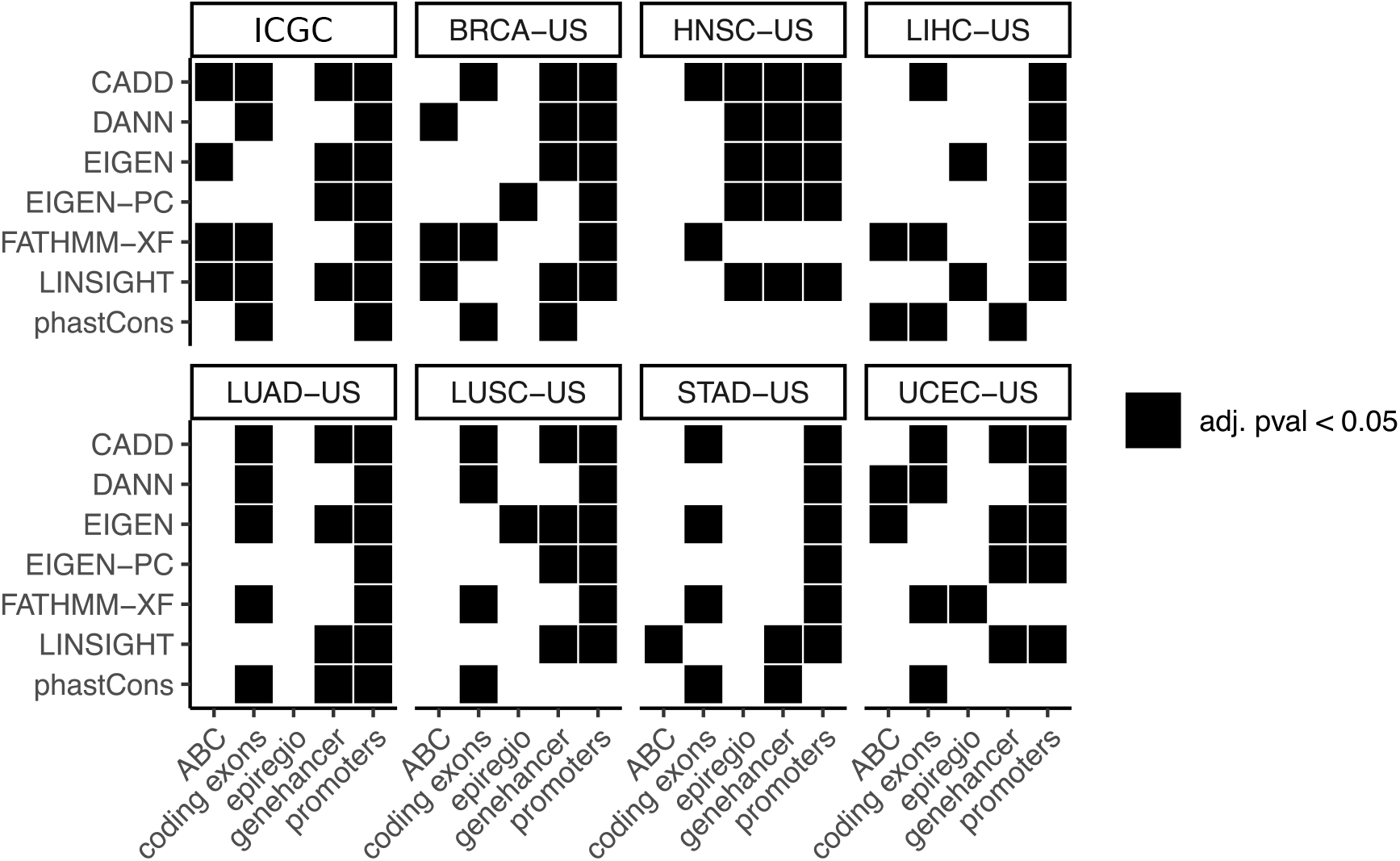
Mutations in cancer genes show significantly higher FI scores. Comparison of cancer versus non-cancer FI scores using different association (x-axis) and scoring (y-axis) methods. Black squares represent those combinations for which cancer genes show a significantly higher (adjusted p-value ≤ 0.05) FI score than non-cancer genes. ABC associations were not available for the HNSC-US cohort.

### HiFIseek: a high-performance method to detect the enrichment of high-impact variants in a set of associated genomic regions

The lack of consensus across CRR association and FI scoring methods prompted us to systematically explore all possible CRR score combinations. We hypothesized that genes exhibiting an enrichment of FI CRVs across multiple CRR-score combinations are more likely to contain driver events. To this end, we developed HiFIseek to study the enrichment of FI variants in sets of associated genomic regions. HiFIseek is a Snakemake pipeline that explores all combinations of user-provided association and scoring methods across any number of cohorts. HiFIseek employs two distinct enrichment detection methods to identify genes exhibiting an enrichment of FI variants. The first one is OncodriveFML, which compares the average impact score of a set of observed variants to a permutation-based expected average. The second one, which we refer to as OncodriveFML-analytic, is a new analytical approach to the OncodriveFML framework that utilizes the sum of observed FI scores to make the method less computationally intensive and less sensitive to outliers. Indeed, using the mean as a summary statistic can be more biased by outliers. HiFIseek includes the following modules for further exploration of genes passing a determined enrichment significance threshold: (1) differential expression analysis when expression data is available, and ontology enrichment analyses using gene ontology (GO) (25, 26), the network of cancer genes (27), and the cancer gene census (28).

We first assessed our new analytic framework on protein-coding somatic mutations to determine whether it could recover well-known cancer-related genes. In all cohorts, OncodriveFML-analytic detected well-known cancer genes (Figure 4, Supplementary Figures 21-28). For example, the top 10 genes when studying the ICGC breast cancer cohort with CADD scores were: *TP53, TMC6, PIK3CA, RB1, MAP3K1, NF1, ARID1A, APBA2, PTEN*, and *CDKN2A*, eight of which are well-known cancer-related genes. In general, our analytic framework tended to detect more genes than OncodriveFML. Overall, we observed a small agreement across scoring approaches, except in well-studied cancer-related genes such as *TP53* or *PIK3CA* (29). These results highlight not only the differences across FI scoring approaches but also their convergence in only the most well-known cancer-related genes.

**Fig. 4.**
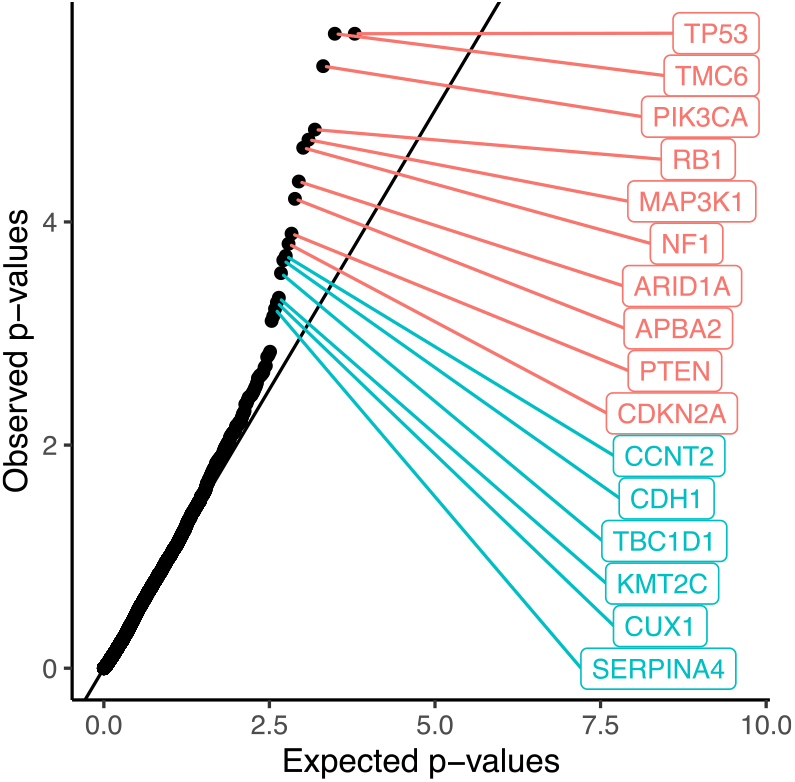
Enrichment of high functional impact mutations in the BASIS cohort detected by OncodriveFML-analytic using CADD scores. Quantile-quantile plot showing the expected (x-axis) versus observed (y-axis) distribution of the log10(p-values) obtained by OncodriveFML-analytic. The diagonal line represents the expected distribution of log_10_ (p-values) if observed p-values follow a uniform distribution. Points deviating from the diagonal towards the upper-left corner indicate significant hits. Genes showing an adjusted p-value *<* 0.1 are labelled in red, while genes with an adjusted p-value between 0.1 and 0.25 are labelled in green.

We next benchmarked OncodriveFML-analytic against OncodriveFML to evaluate their performance in terms of computational requirements. Specifically, we aimed to investigate the required time and memory usage when running OncodriveFML-analytic or OncodriveFML with different numbers of cores, along with gene-region associations and cohorts of increasing size. The dataset used in the bench-marking consisted of: (1) mutations from the ICGC breast cancer (2,277,413 mutations), BRCA-US (1,062,470 mutations), and LIHC-US (717,885 mutations) cohorts; (2) CADD scores; and (3) promoter (1000 bp median size), coding exons (1341 bp median size) and Genehancer (17,890 bp median size) as gene-region associations, which were chosen based on the distribution of region sizes (Supplementary Figure 1). We ran the analyses using 5, 10, 20, 50, or 100 available cores for parallelization and recorded the elapsed time and maximal RAM usage for each analysis. In all analyses, OncodriveFML was run using 1,000,000 permutations. In all cases, OncodriveFML-analytic showed better performance than OncodriveFML both in terms of computation time and required resources (Supplementary Figure 29A-B). Comparison of the results showed a general agreement between the enrichment results of OncodriveFML-analytic and OncodriveFML (Supplementary Figure 29C), with OncodriveFML-analytic tending to yield a more conservative significance estimation than OncodriveFML, particularly for increasing gene-region sizes.

### Exploring the enrichment for high functional impact cis-regulatory variants in eight cancer cohorts

We explored the enrichment of FI somatic mutations in gene CRRs using HiFIseek on all combinations of 8 FI scores (CADD, DANN, EIGEN, EIGEN-PC, FATHMM-XF, LIN-SIGHT, Sei, and phastCons) and 4 gene-CRR associations (promoter regions, ABC, epiregio, and Genehancer). In general, we observed little consensus when comparing all gene-CRR-score combinations within a cohort, showing again the influence of a given association and scoring combination in the analysis (Figure 5, Supplementary Figures 30-36). Gene-hancer regions showed the highest number of genes with a significant enrichment of FI CRVs across all scores and most cohorts. With some exceptions, the majority of genes showing an enrichment of FI CRVs have not been reported to be related to cancer. Recognizing the disparity in results, we focused our attention on those genes found in at least five different combinations of score, region, and/or enrichment method. Gene set enrichment analysis using gene ontology (25, 26), disease ontology (30), or the network of cancer genes (27) did not yield any terms that could be related to cancer. Nevertheless, studying the ICGC breast cancer cohort with HiFIseek resulted in 67 genes showing an enrichment of FI CRVs in at least five combinations, 11 of which have been previously reported to be related to cancer (Figure 5). Among the detected genes, CBFB showed the strongest agreement across FI scores, associations, and enrichment detection methods. This gene has been previously reported to have a tumor suppressor role in breast cancer (31). Another example arose when comparing results across the ICGC and TCGA breast cancer cohorts; we observed that RASGEF1C showed enrichment of FI CRVs in both. This protein is a guanine nucleotide exchange factor, an important molecular function in the activation of Ras signaling (32).

**Fig. 5.**
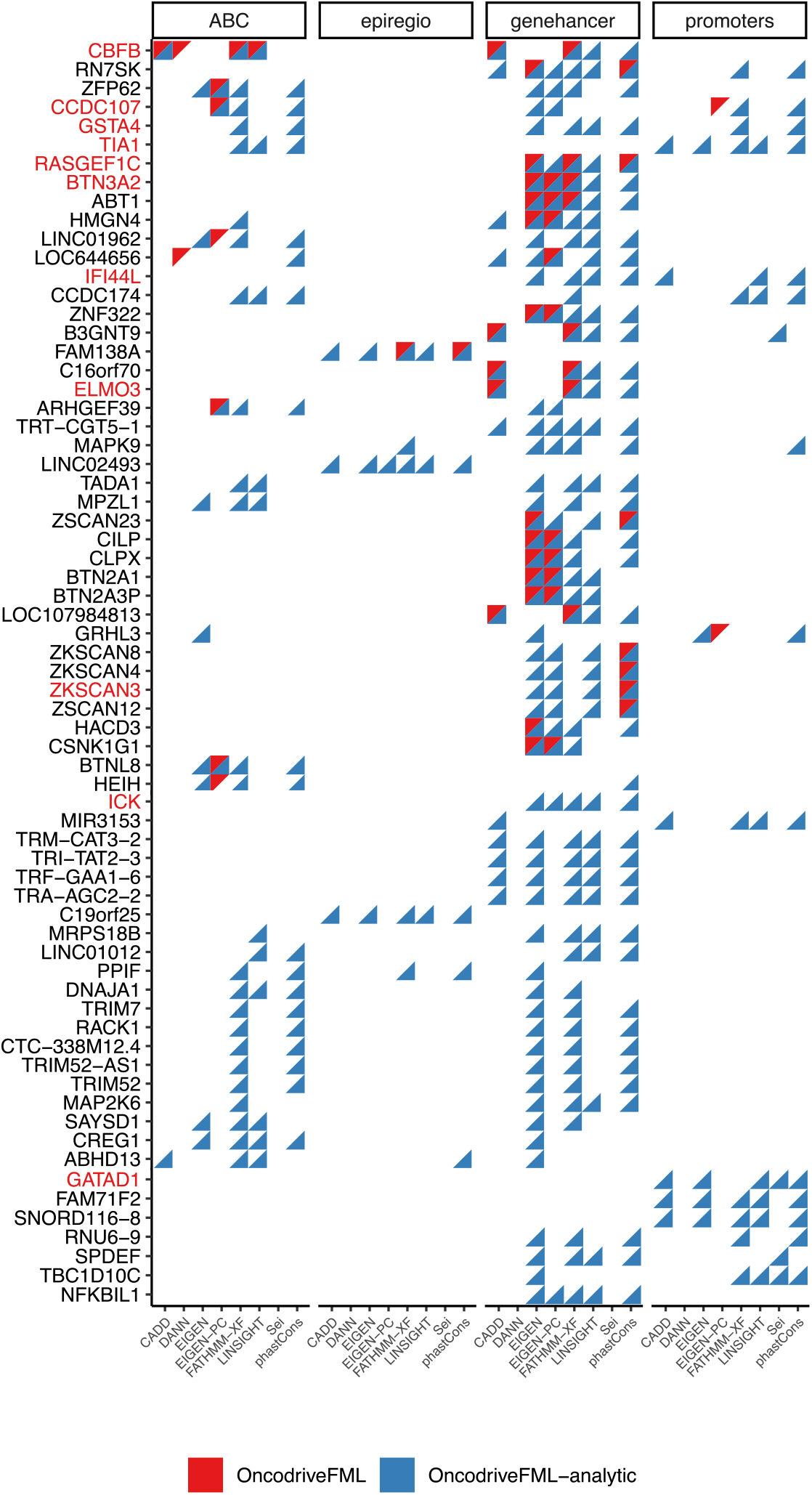
Detected genes across CRR-score combinations in the ICGC breast cancer cohort. Genes (y-axis) showing an enrichment (adjusted-pval ≤ 0.05) of FI CRVs in different score (x-axis) and association (facet) combinations detected by either OncodriveFML (red triangles) or OncodriveFML-analytic (blue triangles). Only genes detected in at least five or more CRR-score combinations and/or enrichment method are shown. Gene names in red are listed as cancer genes.

Given the role of CRRs in transcriptional regulation, we hypothesied that genes with an enrichment of FI CRVs may show changes in their transcriptional levels. To test this, we performed a differential expression analysis for all genes detected by HiFIseek, grouping patients according to the mutation status of each gene’s CRRs. Overall, the majority of genes showing an enrichment of FI CRVs did not show changes in their expression. For example, we did not observe significant changes in the expression levels of CBFB or RAS-GEF1C, where the latter showed a higher but not significant level of expression (Supplementary Figure 37).

### Breast cancer risk SNPs show an enrichment for high impact non-protein-coding variants in known breast cancer related genes

We next applied HiFIseek on a set of SNPs associated with increased breast cancer risk to search for genes with an enrichment of FI SNPs in their CRRs. We ran HiFIseek using promoter and ABC CRRs together with CADD, FATHMM-XF, and SEI scores. Focusing on the genes showing enrichment of FI CRVs in at least two CRR-score combinations resulted in 23 genes, 3 of which have been previously associated with cancer. Given the limited overlap, we examined each combination in more detail. We observed that FATHMM-XF scores combined with ABC CRRs resulted in an enrichment of genes associated with pan-gynecological and breast cancer using the network of cancer genes ontology (Supplementary Figure 39). Based on this, we decided further to explore this combination for potential SNPs of interest. We identified 99 genes exhibiting enrichment of FI SNPs (adjusted p-value ≤ 0.05), including 18 that have been previously reported to be associated with cancer. We identified *BRCA1* among the detected cancer-related genes, a tumor suppressor gene whose loss of function is strongly associated with an increased risk of breast cancer. Given this established role, we next explored the SNPs falling within the CRRs of *BRCA1*. First, we plotted the observed SNP scores in the context of all possible changes in the considered regions to pinpoint those contributing the most to the enrichment (Supplementary Figure 38). In the case of *BRCA1*, SNPs were observed to be at both extremes of the score distribution. Based on this, we further explored the *BRCA1* SNPs showing a FATHMM-XF score ≥ 0.75. We used the UCSC genome browser (33) to study the genomic context surrounding the selected SNPs (Figure 6). Most high-scoring SNPs were located at a genomic region showing active histone marks in HMEC and MCF7 cells. The inclusion of MCF7 cap analysis of gene expression (CAGE) data from FANTOM5 (34) revealed the presence of a lncRNA overlapping with the selected SNPs. Additionally, some of the detected SNPs have been previously reported by FANTOM5 as eQTLs for *BRCA1*.

**Fig. 6.**
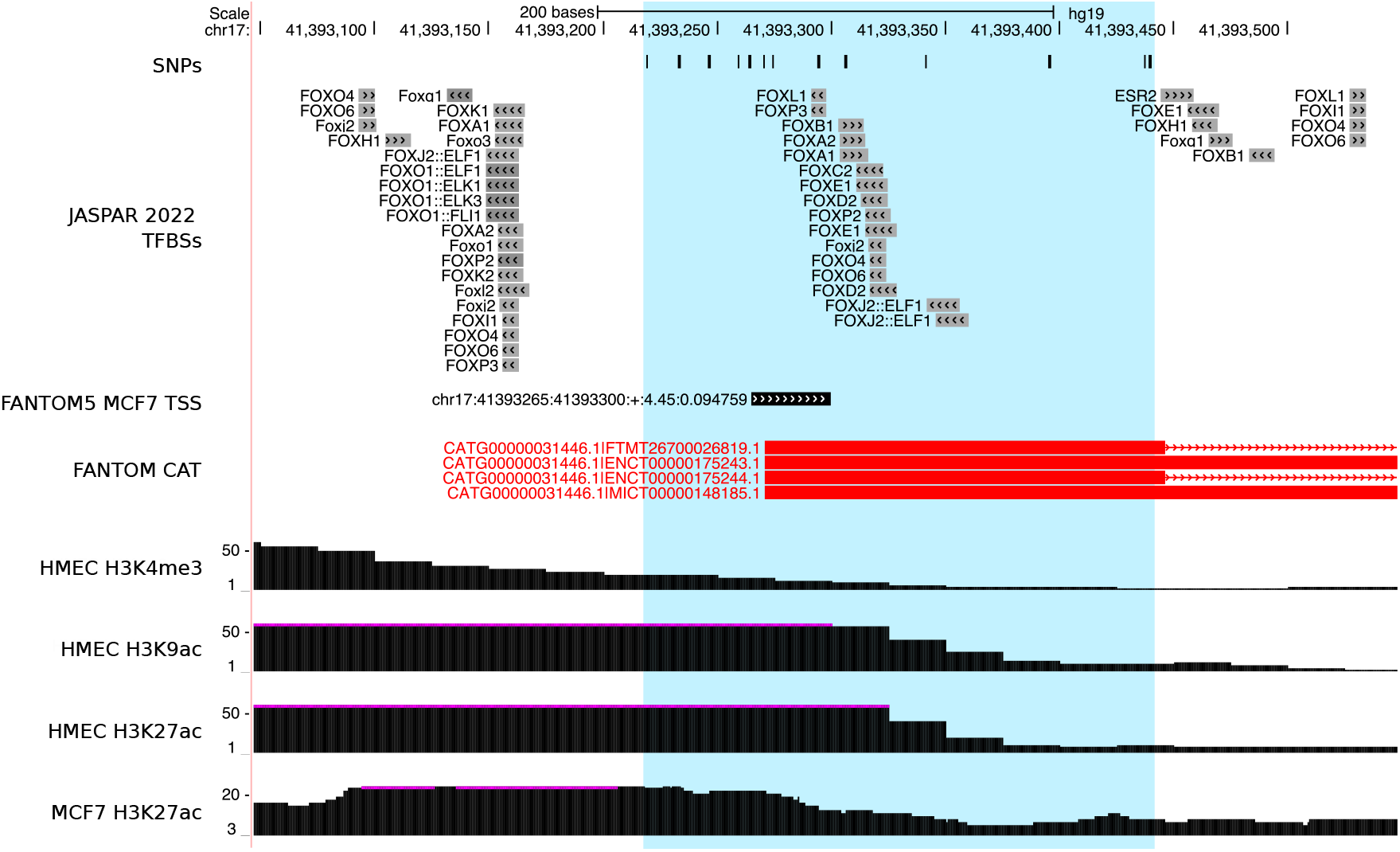
*BRCA1* shows high functional impact SNPs in ABC-derived CRRs. Genomic context around the high FI score SNPs in one of the *BRCA1* CRRs. From top to bottom, tracks show: observed SNPs, JASPAR 2022 TFBSs showing a score higher than 300, FANTOM5 TSS predictions in MCF7 cells, FANTOM CAT transcript predictions, HMEC H3K4me3, H3K9ac and H3K27ac, and MCF7 H3K27Ac.

## Discussion

Searching for potential cis-regulatory driver events is an on-going effort. A significant reason for this is the high complexity of transcriptional regulation. While the basic underlying mechanism is understood, fine-grained, context-specific knowledge of transcriptional regulation remains largely unknown. For example, the same gene may use very different cis-regulatory spaces depending on its cell type and/or condition. Similarly, the same variant may have very different effects in different cellular contexts. Currently, there is a broad range of gene-CRR association and variant impact scoring methods. While they are expected to represent similar mechanisms, we observed a small consensus across association or scoring methods. A likely important factor in this is the input data used in each one of these methods, as well as their underlying assumptions. Nevertheless, the lack of high-quality annotations makes the task of identifying candidate non-protein-coding driver events challenging.

In this work, we describe HiFIseek, a Snakemake pipeline studying the enrichment of high-impact variants in sets of related genomic regions using any number of cohorts, gene-region associations, and FI scoring methods. Additionally, we introduce OncodriveFML-analytic, an analytic approach to the OncodriveFML framework. The main differences between OncodriveFML and our analytic counterpart reside in how the functional impact scores are aggregated and the use of an analytical approach to compute significance. OncodriveFML-analytic demonstrated higher computational performance compared to OncodriveFML, particularly when applied to region associations of increasing sizes. Comparison of OncodriveFML and OncodriveFML-analytic results showed similar results across different analyses, with increasing discrepancy when using larger region associations. We hypothesize that these discrepancies are mainly due to the minimal number of required permutations as the studied regions become larger.

A challenge in detecting gene regions with a higher-than-expected aggregated impact score is striking a balance between a method’s sensitivity and specificity. An important factor in this is the ratio between low and high-scoring variants in the studied regions. For example, a region set showing many low-impact variants and a few high-scoring ones may not be detected, decreasing the method’s specificity. While this could be tackled by pre-filtering very low-scoring mutations, doing so is likely to introduce many false positives and therefore decrease the method’s specificity.

While OncodriveFML-analytic detected many known cancer-related genes when applied to somatic mutations in protein-coding regions, most genes with an enrichment of high-impact CRVs have not been previously recorded to be related to cancer. In CRRs, the amount of disagreement across association and scoring approaches was larger than in the protein-coding space. These results are in agreement with the observed differences between association and FI scoring methods. Additionally, we did not observe significant changes in the transcription levels of the detected genes.

We used HiFIseek to explore gene CRRs using a set of SNPs associated with increased breast cancer risk. While we did not observe a strong consensus across CRR-score combinations, the use of FATHMM scores with ABC CRRs resulted in 18 cancer-related genes with FI SNPs in their CRRs. Among these genes, we identified BRCA1, whose role in breast cancer is well-established. Nevertheless, these results were only supported by one of the explored CRR-score combinations. Given the disagreement across association and scoring approaches, the actual significance of these results is unclear. Likewise, we did not investigate the influence of linkage disequilibrium in our analysis. We further investigated the SNPs in the *BRCA1* due to its known involvement in breast cancer. Manual exploration revealed a subset of SNPs that clustered in a genomic region with signs of activity in both normal and breast cancer cell lines. Given that some of the detected SNPs have been reported as eQTLs for BRCA1, a potential hypothesis is that they increase breast cancer risk by altering *BRCA1* transcription in normal cells. However, the alignment of most high-impact SNPs with an actively transcribed lncRNA in MCF7 cells raises the question of how directly they may influence *BRCA1* transcription. Further work is required to disentangle the involvement of these SNPs in increasing breast cancer risk. Similarly, fine-mapping studies can help identify the most likely causal SNPs.

## Material and Methods

### Cancer data

We used the icgc-get client from the International Cancer Genome Consortium (ICGC) (24) to download mutation, transcription, and copy number alteration data from different TCGA cohorts (23). We used those cohorts for which we could find at least 30 samples with whole genome sequencing (WGS) and RNA-seq data. We downloaded small single nucleotide variants (SNVs) called by MuSE, resulting in 344 samples in total for the following cohorts: BRCA-US (breast invasive carcinoma - 89 donors), LIHC-US (liver hepatocellular carcinoma - 50 donors), UCEC-US (uterine corpus endometrial carcinoma - 48 donors), HNSC-US (head and neck squamous cell carcinoma - 43 donors), LUSC-US (lung squamous cell carcinoma - 42 donors), LUAD-US (lung adenocarcinoma - 37 donors), and STAD-US (stomach adenocarcinoma - 35 donors)(Supplementary Figure 40). Additionally, we obtained 560 WGS samples from the ICGC BASIS breast cancer cohort.

We filtered out hypermutated samples following the approach in (35). Very briefly, we removed samples showing at least 1000 mutations over 1.5 times the interquartile range above the 75th percentile. After filtering, we ended up with the following sample numbers per cohort: BASIS - 520, BRCA-US 85, LIHC-US - 48, UCEC-US - 43, HNSC-US - 40, LUSC-US - 40, LUAD-US - 33, and STAD-US - 32 (Supplementary Figure 40).

RNA-seq raw counts were pre-filtered to remove genes showing 0 reads in *>* 50% of the samples in a cohort. We then normalised raw counts to counts per million and applied a log2 conversion.

### SNPs data

We used observed and imputed genotypes from the Norwegian Breast Cancer Study (NBCS), based on 2,955 individuals, including 2,679 sporadic female breast cancer cases and 276 healthy controls. Samples were genotyped using the iCOGs (36) and OncoArray (37) custom SNP arrays. Imputation was performed as part of the Breast Cancer Association Consortium (38). Participants were recruited between 1975 and 2017 from Oslo University Hospital, Norwegian Radium Hospital, Rikshospitalet, Vestre Viken Hospital Trust, and Akershus University Hospital.

### Gene-region associations

We utilized the following gene-region associations: RefSeq Curated annotations from the UCSC Table Browser, ABC-derived associations, Epiregio, GeneHancer, and superenhancers from SEdb.

The RefSeq curated set of annotations was downloaded on March 28 2022 from the UCSC Table Browser (ncbiRefSeqCurated table) and was used to obtain the following annotations for each gene: (1) 1000bp sequences upstream of their TSS, which we refer to as promoter regions; (2) 5’ UTRs; (3) coding sequences; (4) intronic regions; and (5) 3’ UTRs. Additionally, we used *bedtools subtract* (version 2.30) to remove any promoter sequences overlapping with neighboring gene bodies, thereby eliminating potential confounding effects. Finally, we translated RefSeq transcript IDs to Entrez gene IDs and used *bedtools merge* to merge overlapping genomic regions corresponding to the same gene, thereby avoiding duplicated genomic windows.

ABC associations were downloaded on Feb 9 2022 from ftp://ftp.broadinstitute.org/outgoing/lincRNA/ABC/AllPredictions.AvgHiC.ABC0.015.minus150.ForABCPaperV3.txt.gz and were further processed to fit the required format for OncodriveFML and HiFIseek. We converted gene symbols to Entrez IDs and subtracted regions overlapping with exonic sequences. Finally, we created cohort-specific ABC region subsets by selecting gene-region associations derived from cell types similar to the affected types in a given cohort. Briefly, breast cancer samples used regions annotated as mammary_epithelial_cell-Roadmap, liver hepatocellular carcinoma samples used regions annotated as liver-ENCODE, lung carcinomas used regions annotated as IMR90-Roadmap, stomach adenocarcinoma used regions annotated under stomach-Roadmap, and uterine corpus endometrial carcinoma used regions annotated as uterus-ENCODE.

Epiregio regions were obtained from https://zenodo.org/record/3758494/files/REMAnnotationModelScore_1.csv.gz on July 12 2022. We selected those regions showing a p-value <= 0.05, converted Ensembl IDs to Entrez IDs, lifted them to hg19 using the UCSC liftOver tool (33), merged overlapping intervals associated to the same gene, and subtracted regions overlapping with exonic sequences.

GeneHancer associations (version 4.9) were obtained on February 21 2019. Processing of these regions included selection of elite enhancers, conversion of gene symbols to Entrez IDs, lifting to hg19, merging overlapping intervals for regions associated with the same gene, and subtracting exonic regions.

### Scoring methods

The following functional impact scoring methods were used in this work: CADD (v1.6, obtained on Mar 5 2021), DANN (obtained on Feb 1 2018), EIGEN (obtained on Feb 2 2018), EIGEN-PC (obtained on Feb 4 2018), FATHMM-XF (coding and non-coding) (obtained on May 30 2022 and Feb 23 2018, respectively), LINSIGHT (obtained on Feb 3 2018), Sei (obtained on Aug 3 2022), and phast-Cons100way (obtained on Jul 18 2022).

### Cancer genes

We built a list of cancer genes by including all unique instances of genes being in any of the three following resources: (1) intogen; (2) network of cancer genes; and (3) cancer gene census. Intogen genes were obtained from https://www.intogen.org/download?file=IntOGen-Drivers-20230531.zip on the 24th Aug 2021. Cancer genes from the network of cancer genes were obtained from http://network-cancer-genes.org/download.php on 18th Oct 2023. Finally, tier 1 and 2 cancer genes from the Cancer Gene Census (v98) were obtained from https://cancer.sanger.ac.uk/cosmic/download on October 18, 2023. The resulting list of cancer genes is used throughout this work to study the number of cancer genes in the list of genes showing an enrichment of FI mutations in their associated regions.

### Descriptive statistics

#### Number of mutations by region type

We intersected the RefSeq annotations (see the Gene-region associations section in Methods) with the filtered mutation sets to determine the number of mutations mapping to each type of region. For each cohort, we used *bedtools intersect* and the *uniq* command to determine the number of unique mutations that intersected with each region set in the RefSeq annotations. Additionally, we annotated as “Other non-coding” all mutations that did not overlap with any of the RefSeq-annotated regions.

#### Mutation rates in cancer versus non-cancer genes

For each cohort and gene-CRR association, we computed mutation rates as the number of nucleotides showing at least one mutation in a gene’s CRRs divided by the total number of nucleotides in the gene’s CRRs. Genes were separated into cancer and non-cancer genes using annotations from the Cancer Gene Census (28). Plots comparing the mutation rates were generated with R (39) using the ggplot2 (40) and ggpubr (41) packages.

#### Distribution of functional impact scores in cancer versus non-cancer genes

We obtained the scores for the mutations falling within each gene’s CRRs for each cohort, gene-region association, and mutation impact scoring method. Genes were then categorised as cancer and non-cancer genes using annotations from the Cancer Gene Census (28). Plots comparing the distribution of impact scores in cancer versus non-cancer genes were generated with R (39) using the ggplot2 (40) and ggpubr (41) packages.

#### Gene-CRR sizes and overlap

For each gene, we obtained its respective CRRs according to ABC, epiregio, Genehancer, and promoters. We also introduced intronic regions to explore the amount of gene CRRs predicted to fall within introns. We next computed the number of nucleotides in the entire space of a gene’s CRRs for each association type and plotted the distribution of gene-CRR sizes using the R ggplot2 (40) and ggridges (42) packages.

Gene-CRR overlaps across association methods were computed using bedtools intersect. More specifically, we computed for each gene the amount of intersecting nucleotides between all possible CRR pairwise comparisons. Due to the asymmetry of genomic region intersection, the proportion of intersecting nucleotides was computed in relation to one of the CRR methods in the comparison. For example, the overlap proportion of nucleotides in association method *A* intersecting with association method *B* for gene *g* was computed as:

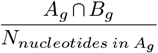

where *A*_*g*_ ∩ *B*_*g*_ is the number of nucleotides in association method *A* intersecting with association method *B* for gene *g* and *N*_*nucleotides in A*_*g* is the amount of nucleotides in association method *A* for gene *g*. Finally, we computed for each pairwise comparison the median overlap across all genes and plotted them using the R ggplot2 package.

### Distribution of impact scores and computation of score agreement

For each scoring method, we recorded the number of times each value was observed and plotted them using the R ggplot2 package. Agreement across scoring approaches was done by randomly sampling 10,000 distinct mutations from the all TCGA cohorts and obtaining their functional impact scores for all considered scoring methods. Finally, we plotted all possible scoring method pairwise comparisons using the R ggplot2 and GGally (43) packages.

### HiFIseek

HiFIseek is a Snakemake-based pipeline computing the enrichment of functional mutations in any number of cohorts, region association, and score combinations. Enrichment is computed through two different enrichment methods: (1) OncodriveFML, and (2) an analytical approach to the OncodriveFML framework using the sum of the observed impact scores.

HiFIseek takes as inputs one or more sets of associated genomic regions (e.g. CRRs for each gene, or coding sequences for a gene), precomputed functional impact scores, and cohorts. The results of the pipeline are output in a Markdown document to facilitate the exploration of all combinations.

The HiFIseek code can be found at https://bitbucket.org/CBGR/hifiseek.

### Enrichment computation with OncodriveFML-analytic

As a first step, OncodriveFML-analytic takes for each region set the mutations intersecting with it and sums their functional impact scores. As a parameter, a user can specify the minimal number of mutations a set of regions must show to be considered in the analysis. Next, it computes the distribution of all possible sums of scores for mutation sets of the same size while taking into consideration all possible mutations in the studied region set and the mutational signature of the studied samples. First, the probability of observing each score is modeled through a multinomial distribution where each class is a functional impact score and the probability of observing it is computed considering: (1) the amount of times it is observed, and (2) the mutational signature of a given cohort. We use the bgsignature python package to compute the signature for a given cancer cohort. For each region set, we next obtain the signature probability and normalise it by dividing it by the sum of probabilities for all mutations in the studied region. Normalised probabilities are then aggregated by score value. We then model the distribution of score sums for an observed mutation set of size *M* as a normal distribution *N* (*μ, σ*), where:

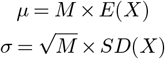

where *E*(*X*) and *SD*(*X*) are the mean and standard deviation of the multinomial distribution for the studied region set. Significance for the observed combined score is then calculated by computing the probability of observing an equal or higher aggregated score than the observed score. Finally, multiple testing correction is applied to all the studied region sets using the Benjamini-Hochberg method.

### Computational benchmark

We analysed the set of filtered somatic mutations in the BRCA-US cohort using CADD scores and the following gene-region associations: promoters, coding exons, and introns. The regions were selected based on the distribution of region sizes to explore the computational impact of exploring larger genomic windows. We ran each analysis using different numbers of cores to observe how the method scales as more resources can be used in parallel. All OncodriveFML analyses were run using 1,000,000 permutations. For each analysis, we recorded the total elapsed time and aggregated RAM usage across all processes. Plots comparing the results between OncodriveFML and OncodriveFML-analytic show the raw p-values reported by either OncodriveFML or OncodriveFML-analytic. Plots were generated using the ggplot2 R package.

## Acknowledgements

We thank the members of the CBGR and Kuijjer groups for their valuable feedback during discussions. We thank the NCMM IT team for their IT support and Ingrid Kjelsvik for administrative support.

