## Supplementary for "HiFIseek: gene-specific enrichment of high-impact mutations in associated genomic regions"

### Supplementary Material

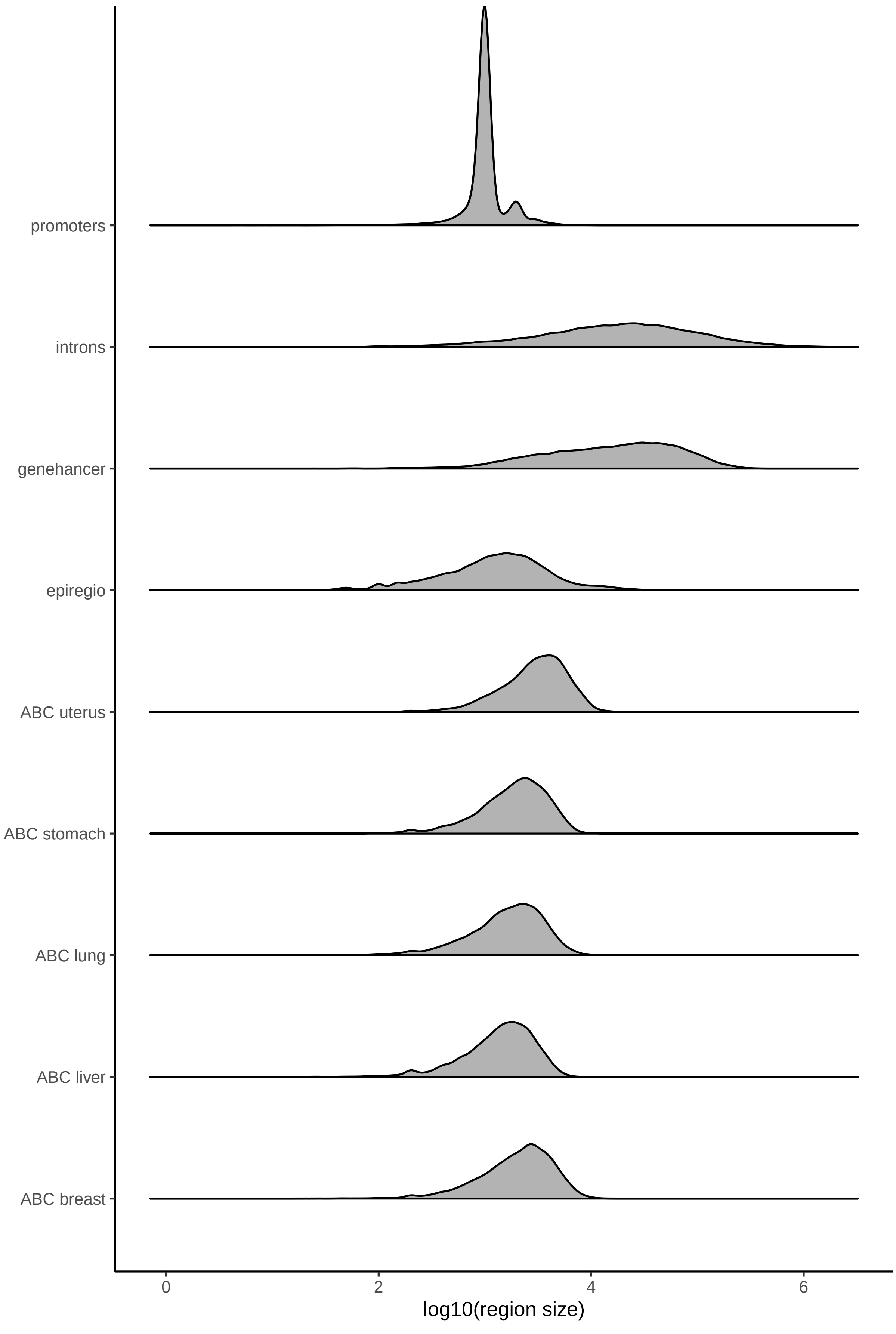


**Supplementary Figure 1: Distribution of gene-region sizes across association methods.** Density plots showing the $\log_{10}$(aggregated region size) for each gene (x-axis) according to different gene-CRR association methods (y-axis).

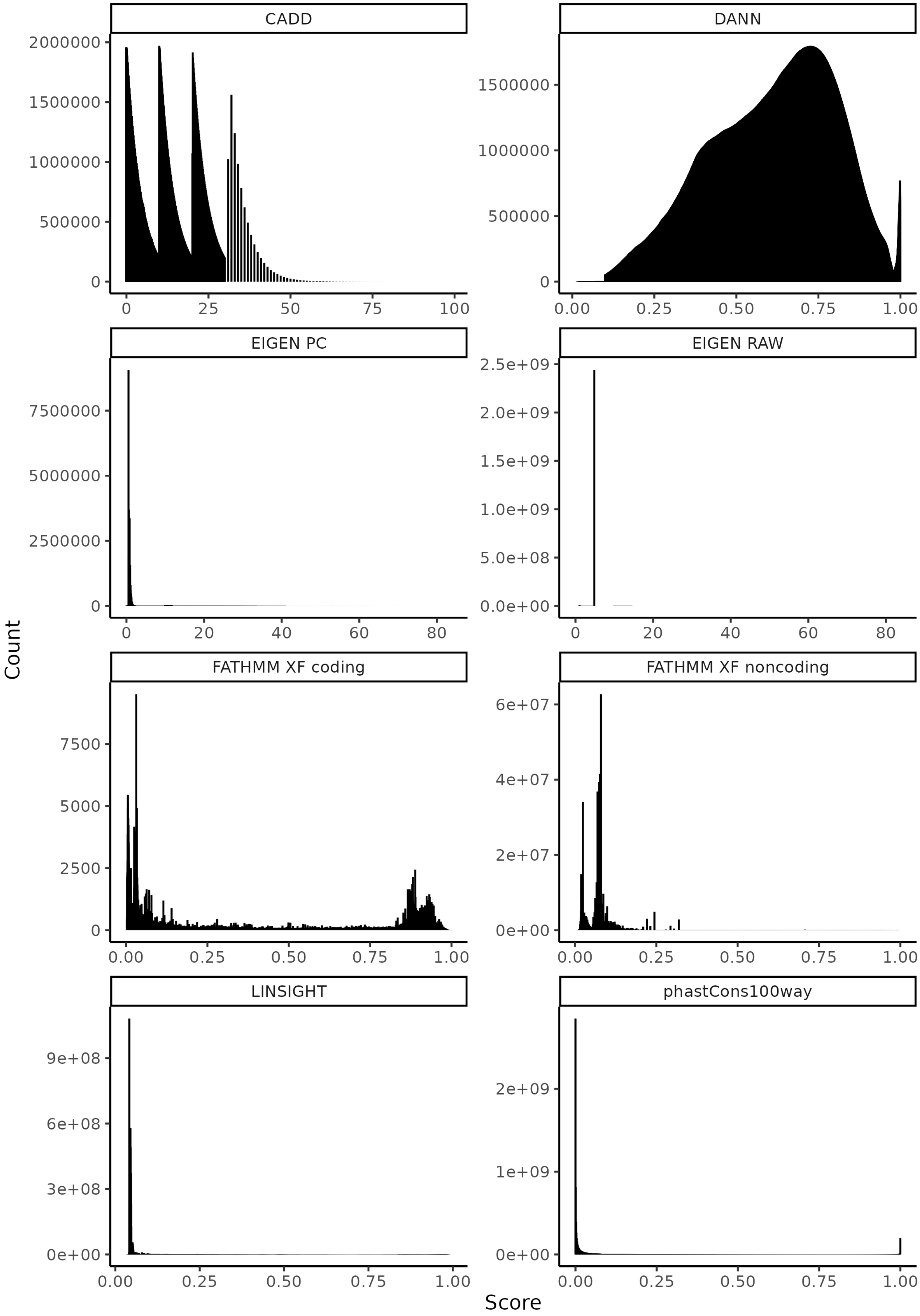


**Supplementary Figure 2: Mutation impact scores show different distributions.** Barplots showing the frequency (y-axis) of each impact score (x-axis) separated by scoring methods.

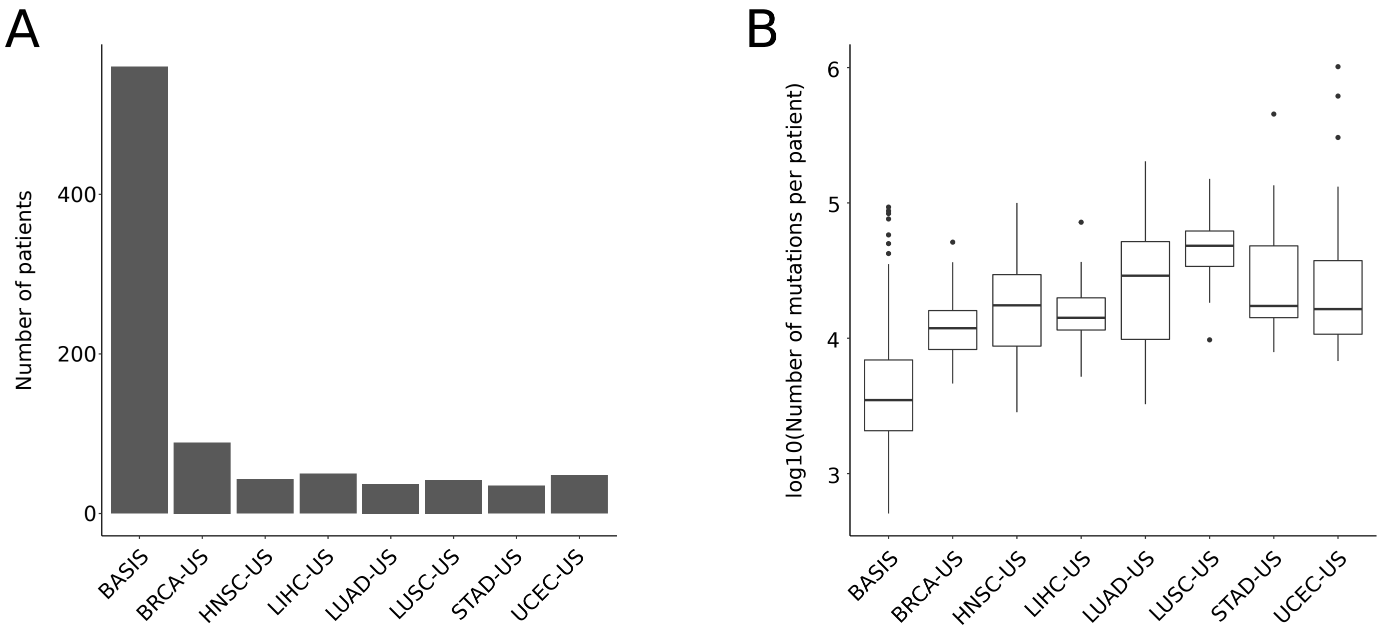


**Supplementary Figure 3: Descriptive statistics of the used cohorts.** (A) Number of patients (y-axis) per cohort (x-axis). (B) $\log_{10}$(Number of mutations per patient) (y-axis) per cohort (x-axis).

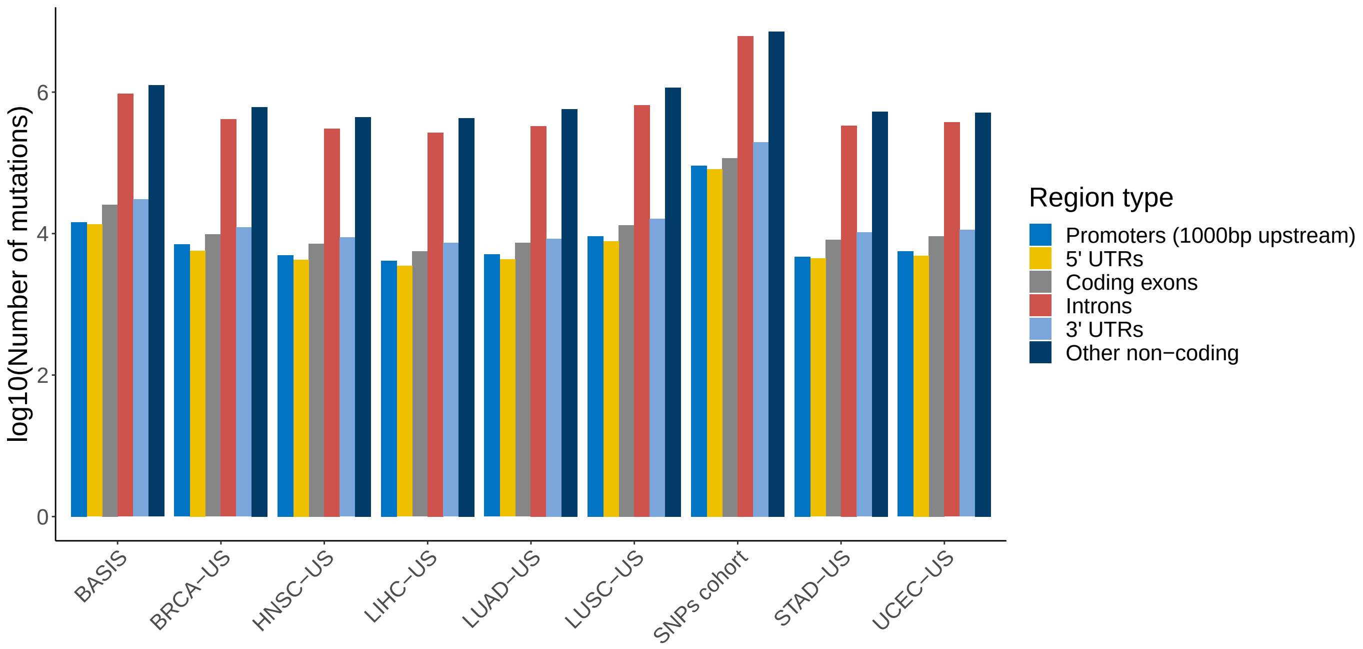


**Supplementary Figure 4: Number of unique mutations in different types of genomic regions.** $\mathrm{Log}_{10}$(number of unique mutations) (y-axis) falling in promoters, 5’ UTRs, coding exons, introns, 3’ UTRs, and other non-coding regions by cancer cohort (x-axis).

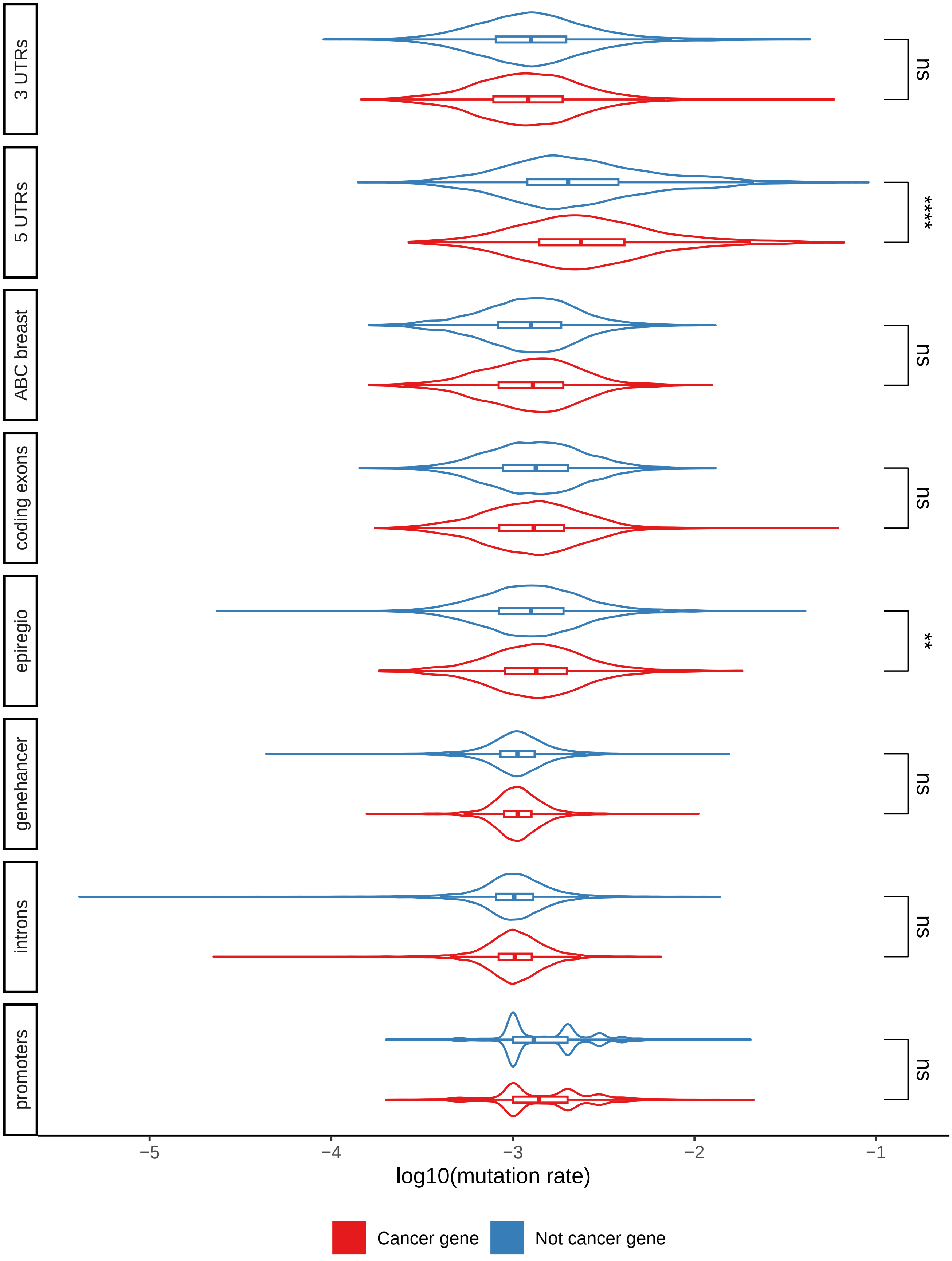


**Supplementary Figure 5: Mutation rates per region type for the BASIS cohort.** $\mathrm{Log}_{10}$(mutation rates) for cancer (red) and non-cancer (blue) genes stratified by gene-region association type. Significance labels indicate the p-value for the one-sided Wilcoxon test checking whether the $\log_{10}$(mutation rate) for the cancer genes $>$ $\log_{10}$(mutation rate) for the non-cancer genes. Significance labels indicate: ns - $p > 0.05$; $* : p \leq 0.05$; $** : p \leq 0.01$; $*** : p \leq 0.001$; $**** : p \leq 0.0001$.

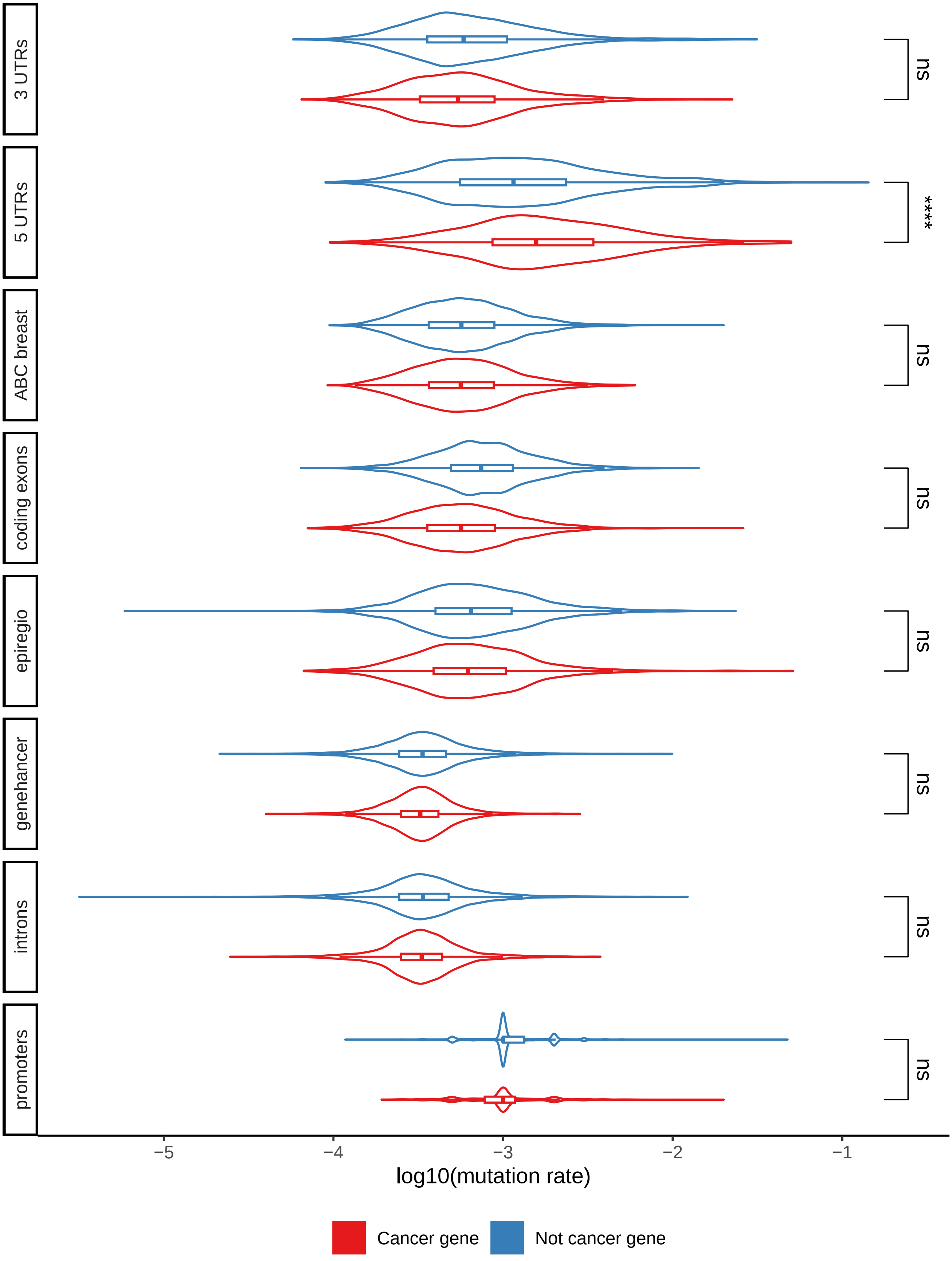


**Supplementary Figure 6: Mutation rates per region type for the BRCA-US cohort.** $\text{Log}_{10}$(mutation rates) for cancer (red) and non-cancer (blue) genes stratified by gene-region association type. Significance labels indicate the p-value for the one-sided Wilcoxon test checking whether the $\log_{10}$(mutation rate) for the cancer genes $>$ $\log_{10}$(mutation rate) for the non-cancer genes. Significance labels indicate: ns - $p > 0.05$; $* : p \leq 0.05$; $** : p \leq 0.01$; $*** : p \leq 0.001$; $**** : p \leq 0.0001$.

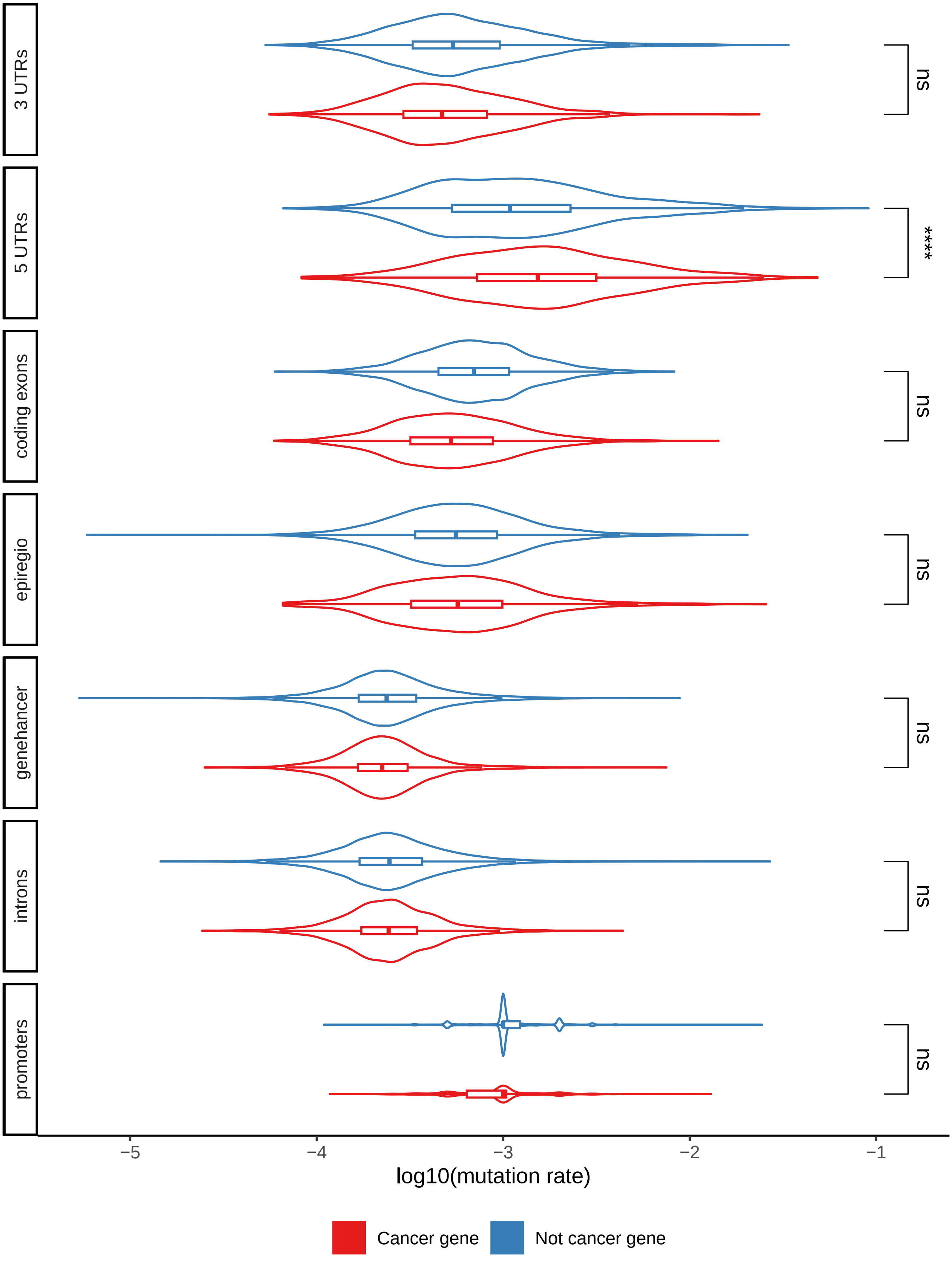


**Supplementary Figure 7: Mutation rates per region type for the HNSC-US cohort.** $\text{Log}_{10}$(mutation rates) for cancer (red) and non-cancer (blue) genes stratified by gene-region association type. Significance labels indicate the p-value for the one-sided Wilcoxon test checking whether the $\log_{10}$(mutation rate) for the cancer genes > $\log_{10}$(mutation rate) for the non-cancer genes. Significance labels indicate: ns - $p > 0.05$; $* : p \leq 0.05$; $** : p \leq 0.01$; $*** : p \leq 0.001$; $**** : p \leq 0.0001$.

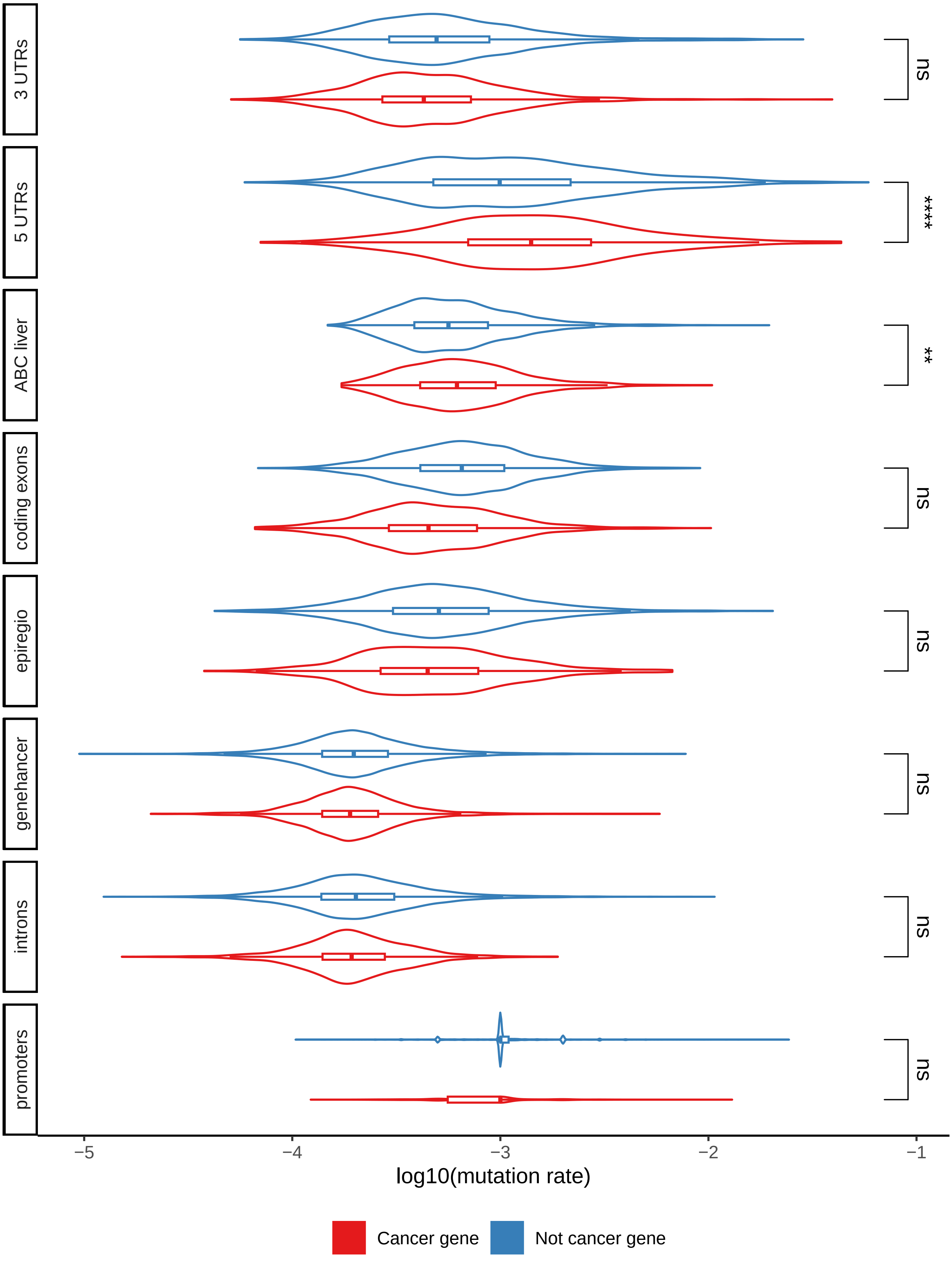


**Supplementary Figure 8: Mutation rates per region type for the LIHC-US cohort.** $\text{Log}_{10}$(mutation rates) for cancer (red) and non-cancer (blue) genes stratified by gene-region association type. Significance labels indicate the p-value for the one-sided Wilcoxon test checking whether the $\log_{10}$(mutation rate) for the cancer genes $>$ $\log_{10}$(mutation rate) for the non-cancer genes. Significance labels indicate: ns - $p > 0.05$; $* : p \leq 0.05$; $** : p \leq 0.01$; $*** : p \leq 0.001$; $**** : p \leq 0.0001$.

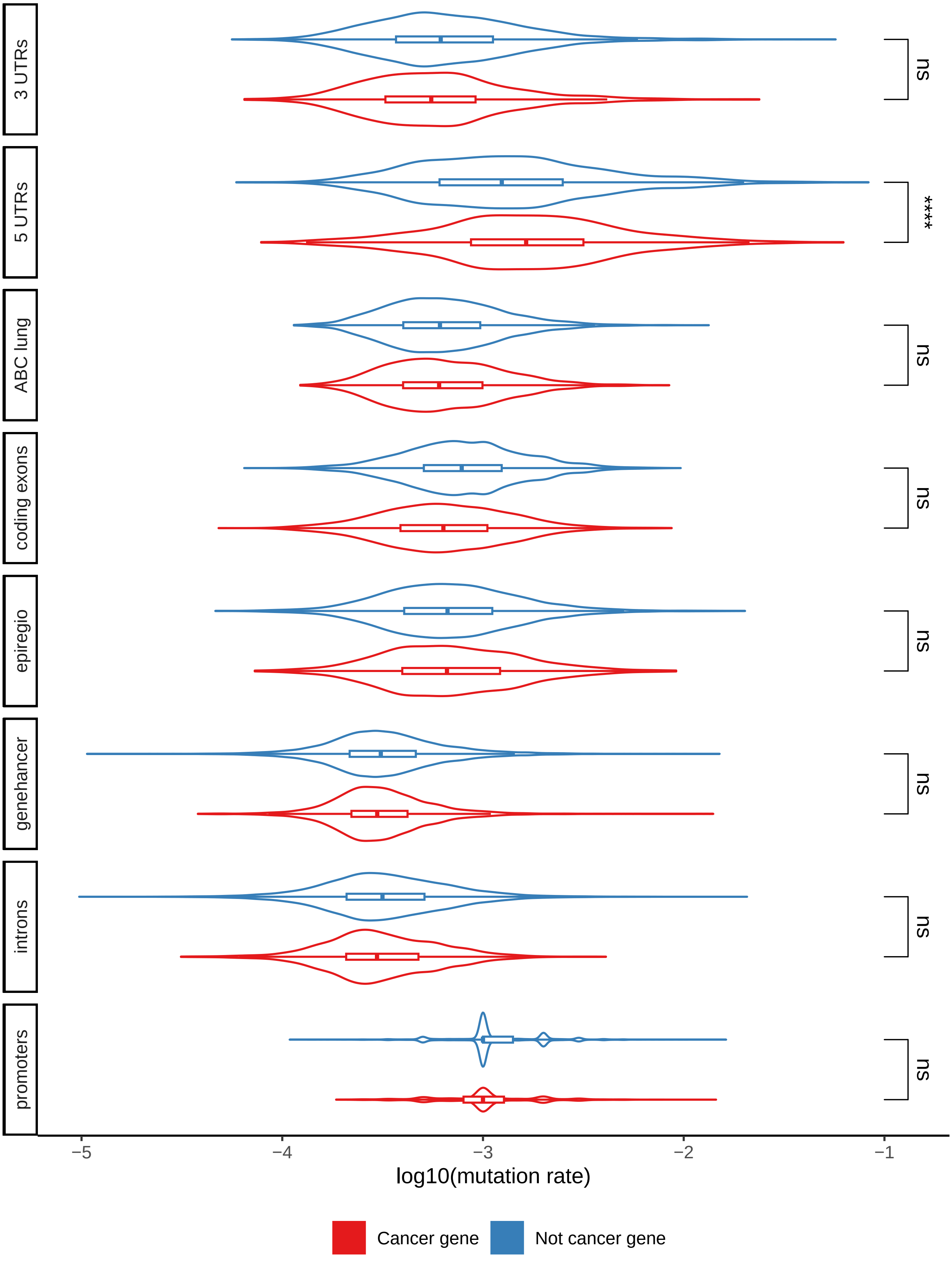


**Supplementary Figure 9: Mutation rates per region type for the LUAD-US cohort.** $\text{Log}_{10}$(mutation rates) for cancer (red) and non-cancer (blue) genes stratified by gene-region association type. Significance labels indicate the p-value for the one-sided Wilcoxon test checking whether the $\log_{10}$(mutation rate) for the cancer genes $>$ $\log_{10}$(mutation rate) for the non-cancer genes. Significance labels indicate: ns - $p > 0.05$; $* : p \leq 0.05$; $** : p \leq 0.01$; $*** : p \leq 0.001$; $**** : p \leq 0.0001$.

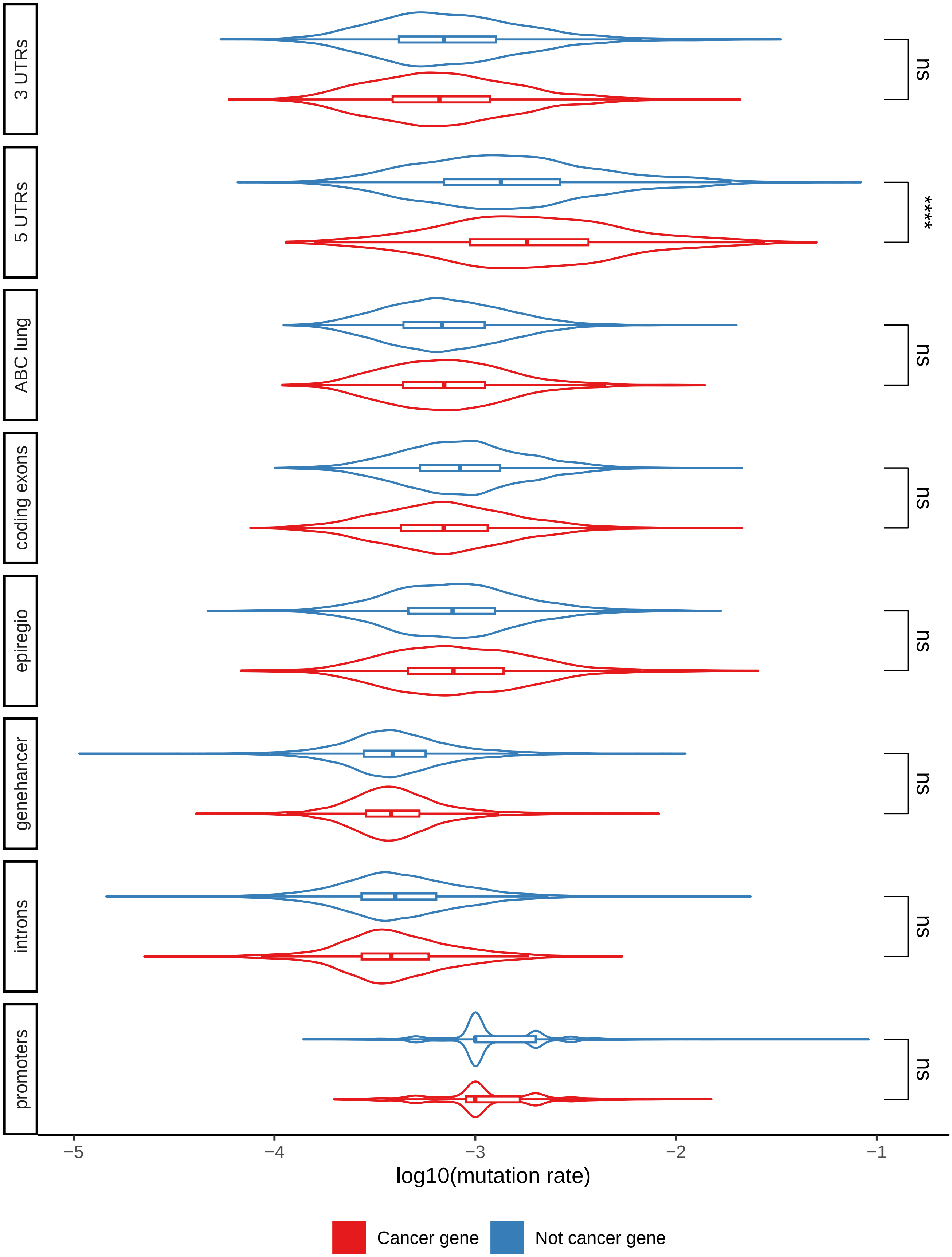


**Supplementary Figure 10: Mutation rates per region type for the LUSC-US cohort.** $\text{Log}_{10}$(mutation rates) for cancer (red) and non-cancer (blue) genes stratified by gene-region association type. Significance labels indicate the p-value for the one-sided Wilcoxon test checking whether the $\log_{10}$(mutation rate) for the cancer genes $>$ $\log_{10}$(mutation rate) for the non-cancer genes. Significance labels indicate: ns - $p > 0.05$; $* : p \leq 0.05$; $** : p \leq 0.01$; $*** : p \leq 0.001$; $**** : p \leq 0.0001$.

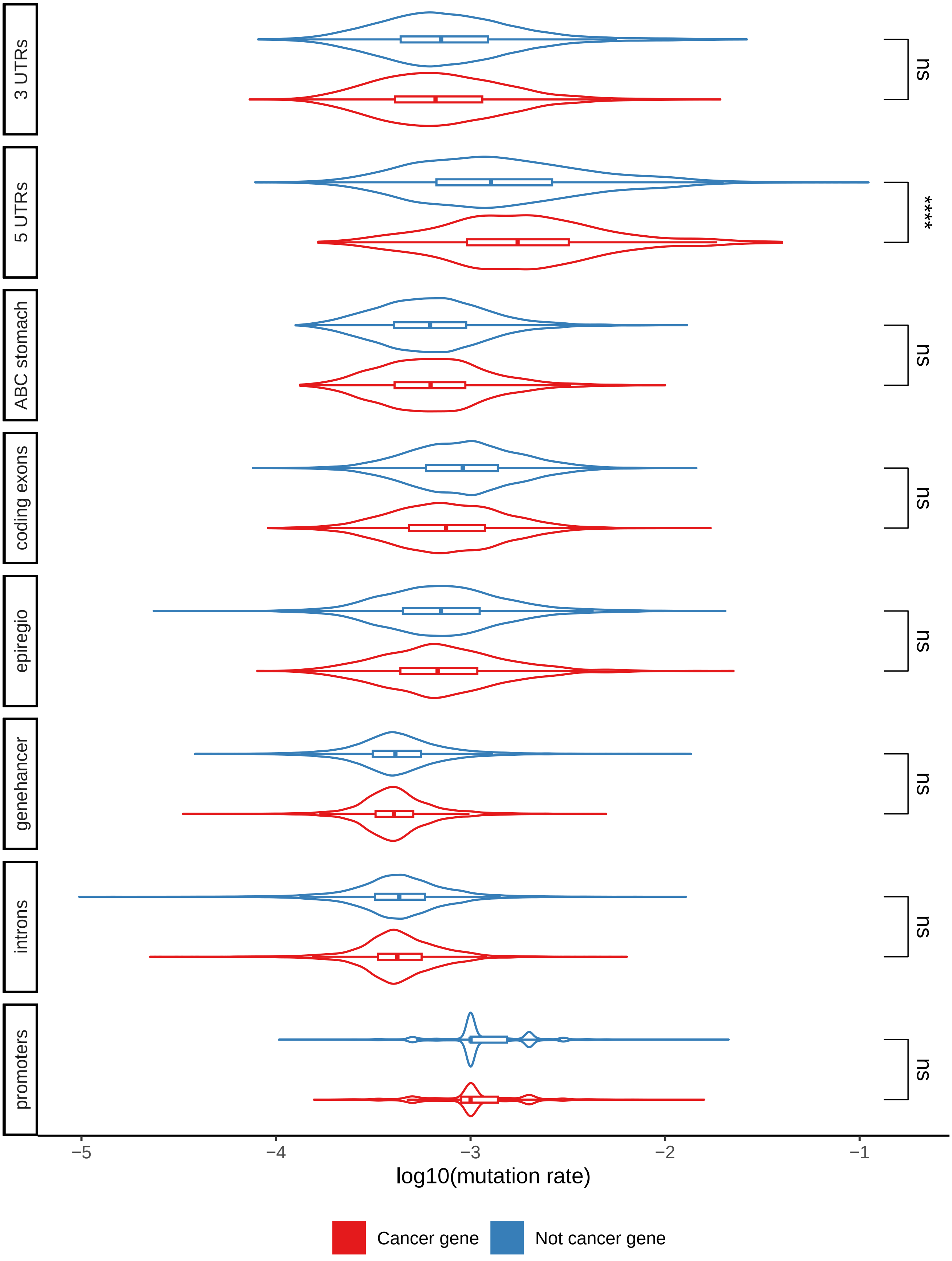


**Supplementary Figure 11: Mutation rates per region type for the STAD-US cohort.** $\text{Log}_{10}$(mutation rates) for cancer (red) and non-cancer (blue) genes stratified by gene-region association type. Significance labels indicate the p-value for the one-sided Wilcoxon test checking whether the $\log_{10}$(mutation rate) for the cancer genes $>$ $\log_{10}$(mutation rate) for the non-cancer genes. Significance labels indicate: ns - $p > 0.05$; $* : p \leq 0.05$; $** : p \leq 0.01$; $*** : p \leq 0.001$; $**** : p \leq 0.0001$.

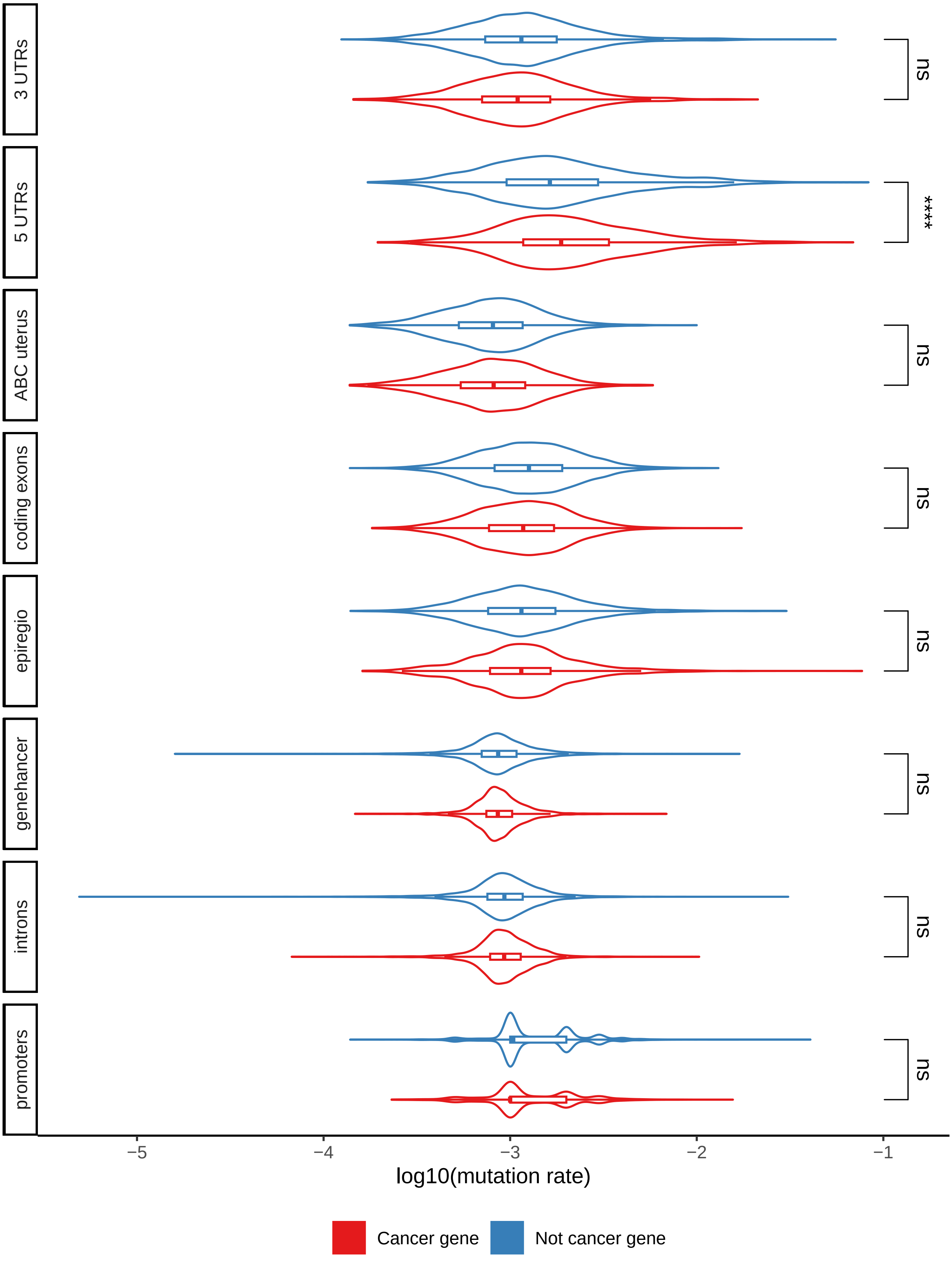


**Supplementary Figure 12: Mutation rates per region type for the UCEC-US cohort.** $\text{Log}_{10}$(mutation rates) for cancer (red) and non-cancer (blue) genes stratified by gene-region association type. Significance labels indicate the p-value for the one-sided Wilcoxon test checking whether the $\log_{10}$(mutation rate) for the cancer genes $>$ $\log_{10}$(mutation rate) for the non-cancer genes. Significance labels indicate: ns - $p > 0.05$; $*$ : $p \leq 0.05$; $**$ : $p \leq 0.01$; $***$ : $p \leq 0.001$; $****$ : $p \leq 0.0001$.

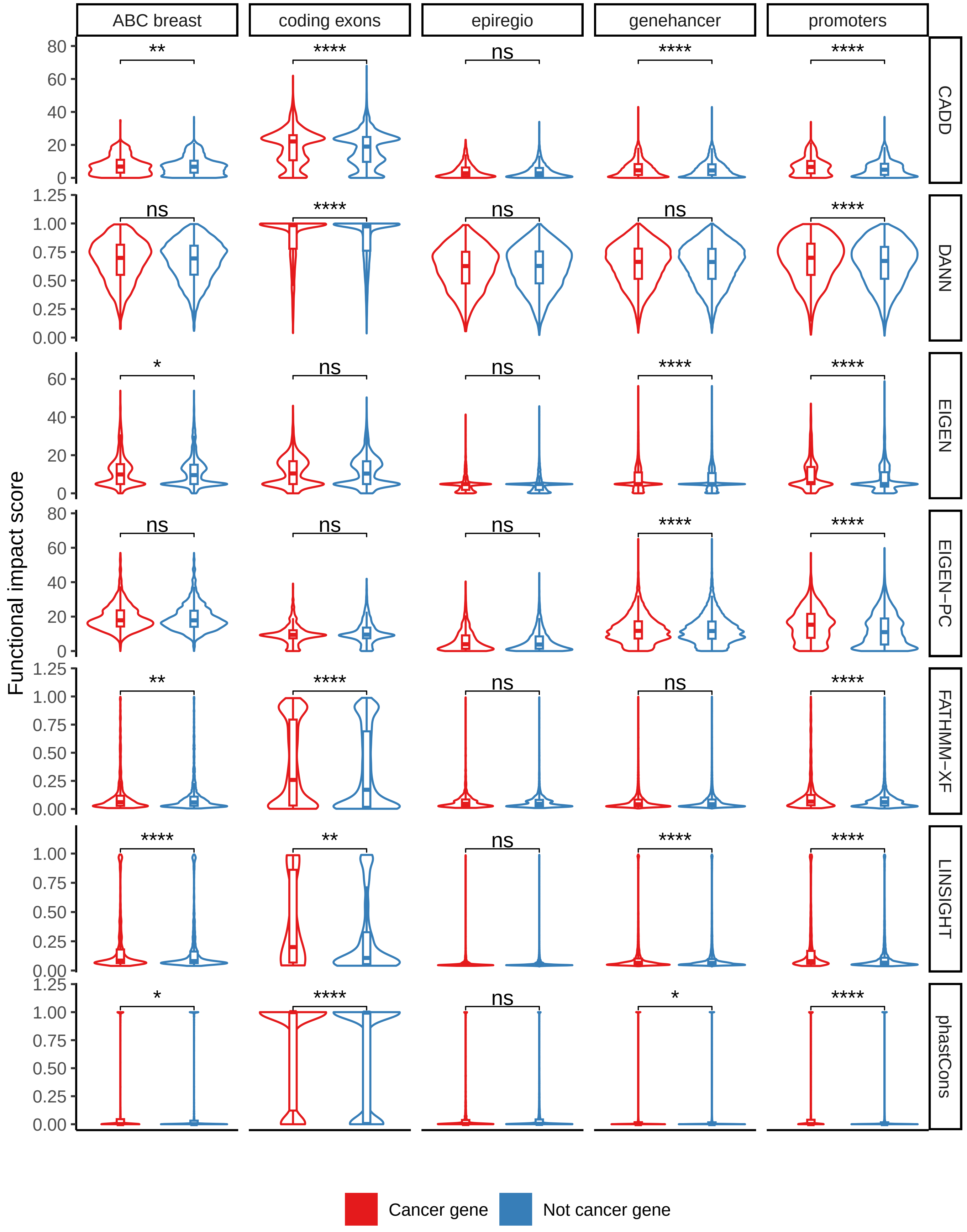


**Supplementary Figure 13: Impact score distributions per region and score type for the observed mutations in the BASIS cohort.** Distribution of functional impact scores for cancer (red) and non-cancer (blue) genes separated by association and scoring type. Significance labels indicate the p-value for the one-sided Wilcoxon test checking whether the $\log_{10}$(mutation rate) for the cancer genes $>$ $\log_{10}$(mutation rate) for the non-cancer genes. Significance labels indicate: ns - $p > 0.05$; $* : p \leq 0.05$; $** : p \leq 0.01$; $*** : p \leq 0.001$; $**** : p \leq 0.0001$.

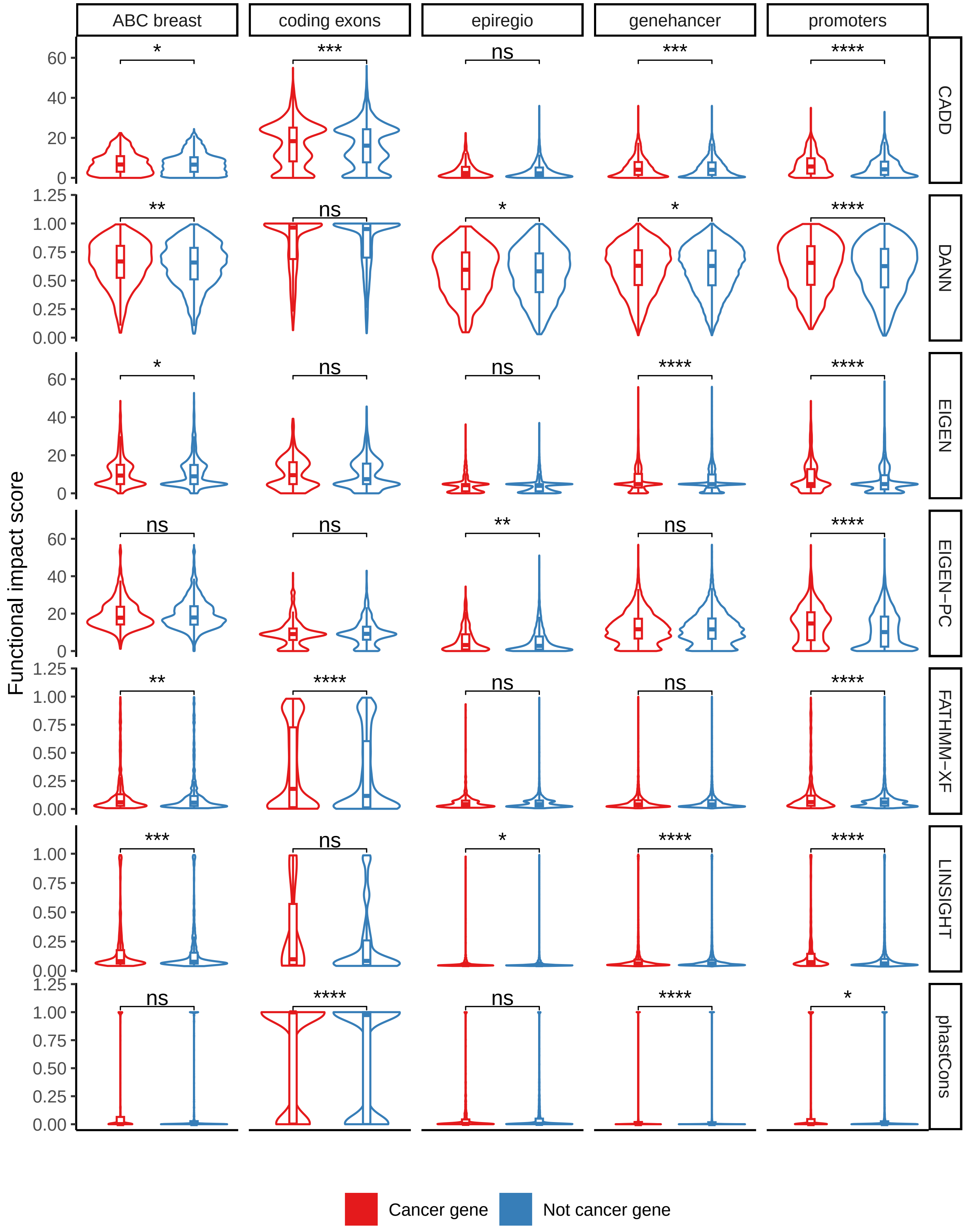


**Supplementary Figure 14: Impact score distributions per region and score type for the observed mutations in the BRCA-US cohort.** Distribution of functional impact scores for cancer (red) and non-cancer (blue) genes separated by association and scoring type. Significance labels indicate the p-value for the one-sided Wilcoxon test checking whether the $\log_{10}$(mutation rate) for the cancer genes $>$ $\log_{10}$(mutation rate) for the non-cancer genes. Significance labels indicate: ns - $p > 0.05$; $* : p \leq 0.05$; $** : p \leq 0.01$; $*** : p \leq 0.001$; $**** : p \leq 0.0001$.

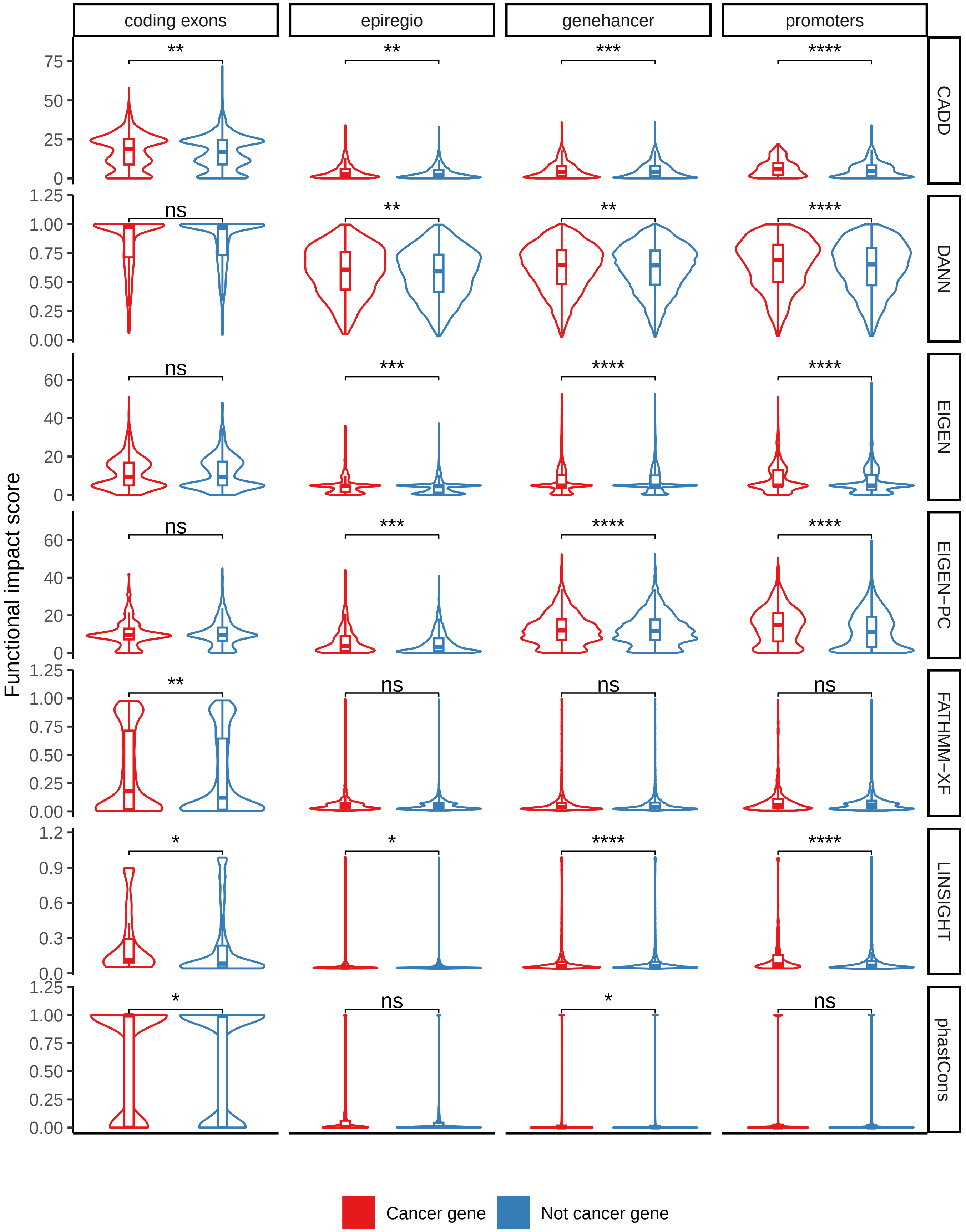


**Supplementary Figure 15: Impact score distributions per region and score type for the observed mutations in the HNSC-US cohort.** Distribution of functional impact scores for cancer (red) and non-cancer (blue) genes separated by association and scoring type. Significance labels indicate the p-value for the one-sided Wilcoxon test checking whether the $\log_{10}$(mutation rate) for the cancer genes $>$ $\log_{10}$(mutation rate) for the non-cancer genes. Significance labels indicate: ns - $p > 0.05$; $* : p \leq 0.05$; $** : p \leq 0.01$; $*** : p \leq 0.001$; $**** : p \leq 0.0001$.

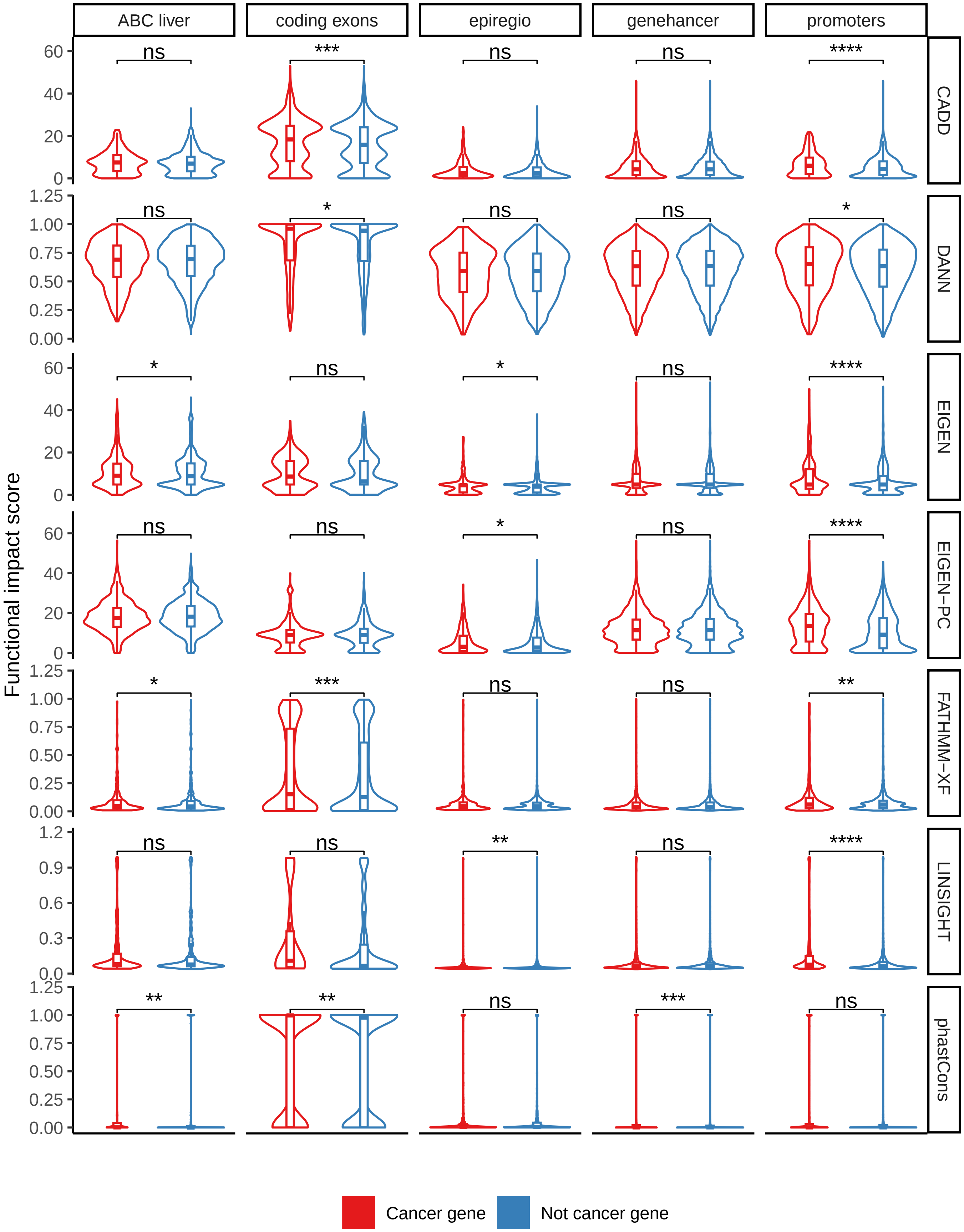


**Supplementary Figure 16: Impact score distributions per region and score type for the observed mutations in the LIHC-US cohort.** Distribution of functional impact scores for cancer (red) and non-cancer (blue) genes separated by association and scoring type. Significance labels indicate the p-value for the one-sided Wilcoxon test checking whether the $\log_{10}$(mutation rate) for the cancer genes $>$ $\log_{10}$(mutation rate) for the non-cancer genes. Significance labels indicate: ns - $p > 0.05$; $* : p \leq 0.05$; $** : p \leq 0.01$; $*** : p \leq 0.001$; $**** : p \leq 0.0001$.

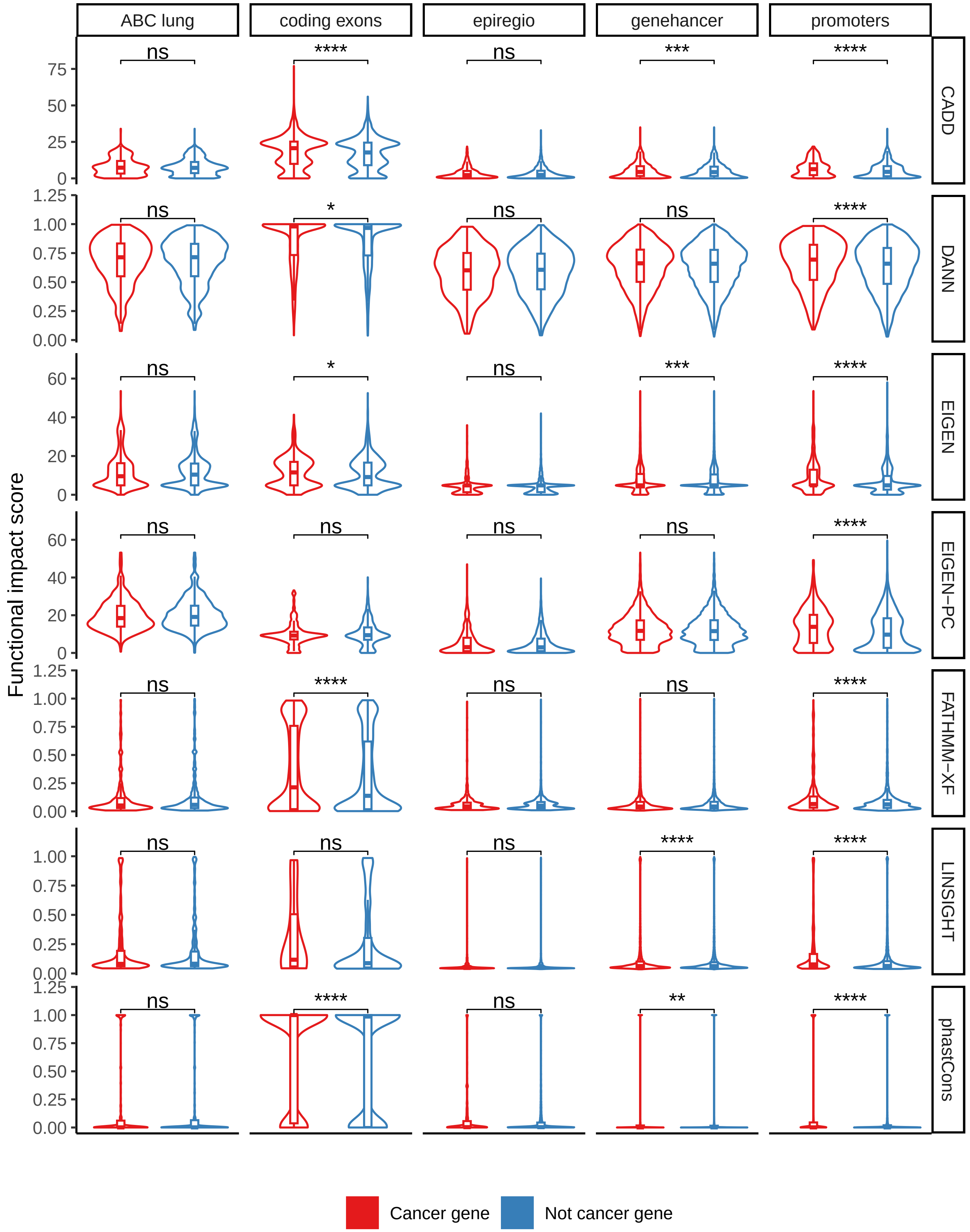


**Supplementary Figure 17: Impact score distributions per region and score type for the observed mutations in the LUAD-US cohort.** Distribution of functional impact scores for cancer (red) and non-cancer (blue) genes separated by association and scoring type. Significance labels indicate the p-value for the one-sided Wilcoxon test checking whether the $\log_{10}$(mutation rate) for the cancer genes $>$ $\log_{10}$(mutation rate) for the non-cancer genes. Significance labels indicate: ns - $p > 0.05$; $* : p \leq 0.05$; $** : p \leq 0.01$; $*** : p \leq 0.001$; $**** : p \leq 0.0001$.

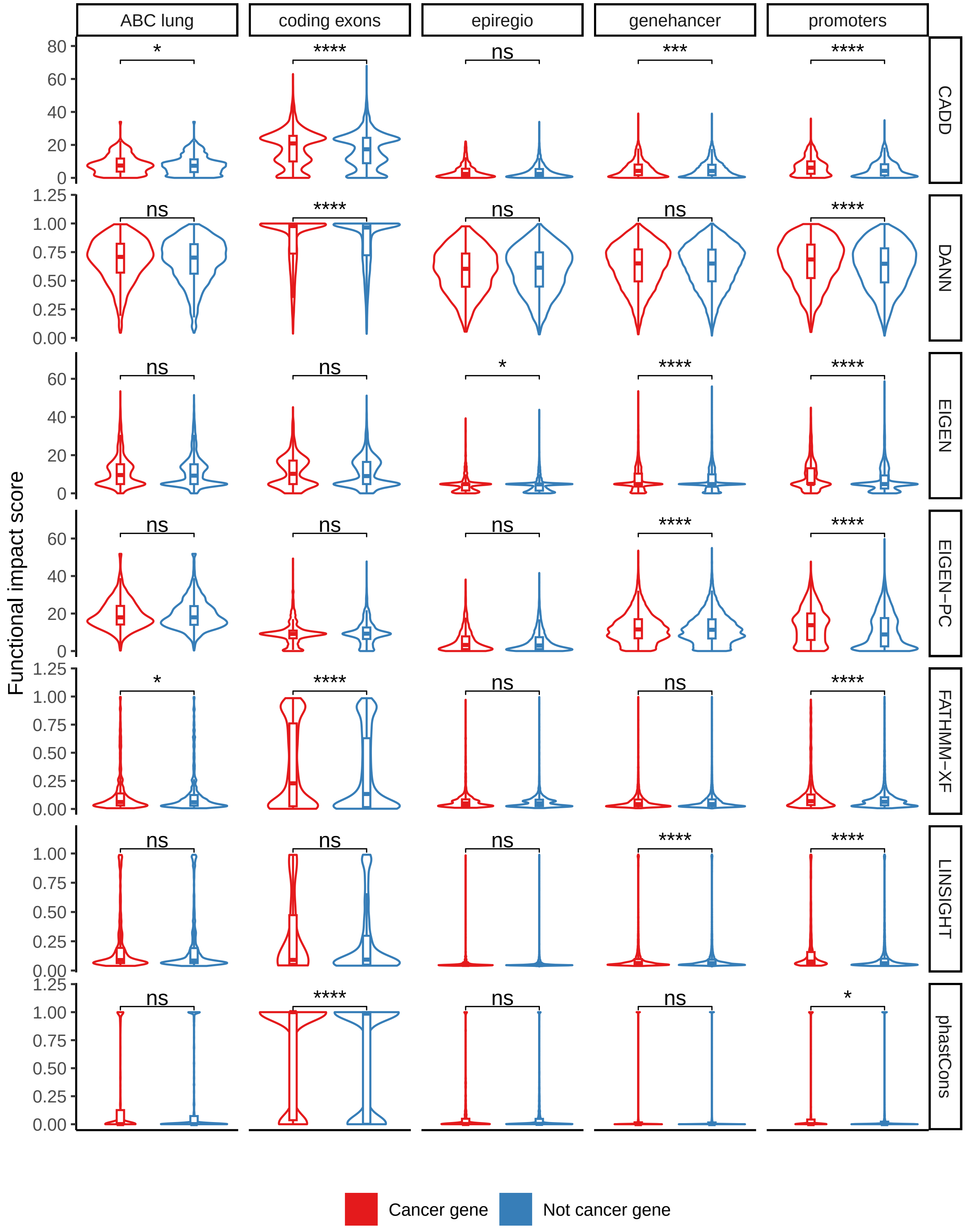


**Supplementary Figure 18: Impact score distributions per region and score type for the observed mutations in the LUSC-US cohort.** Distribution of functional impact scores for cancer (red) and non-cancer (blue) genes separated by association and scoring type. Significance labels indicate the p-value for the one-sided Wilcoxon test checking whether the $\log_{10}$(mutation rate) for the cancer genes $>$ $\log_{10}$(mutation rate) for the non-cancer genes. Significance labels indicate: ns - $p > 0.05$; $* : p \leq 0.05$; $** : p \leq 0.01$; $*** : p \leq 0.001$; $**** : p \leq 0.0001$.

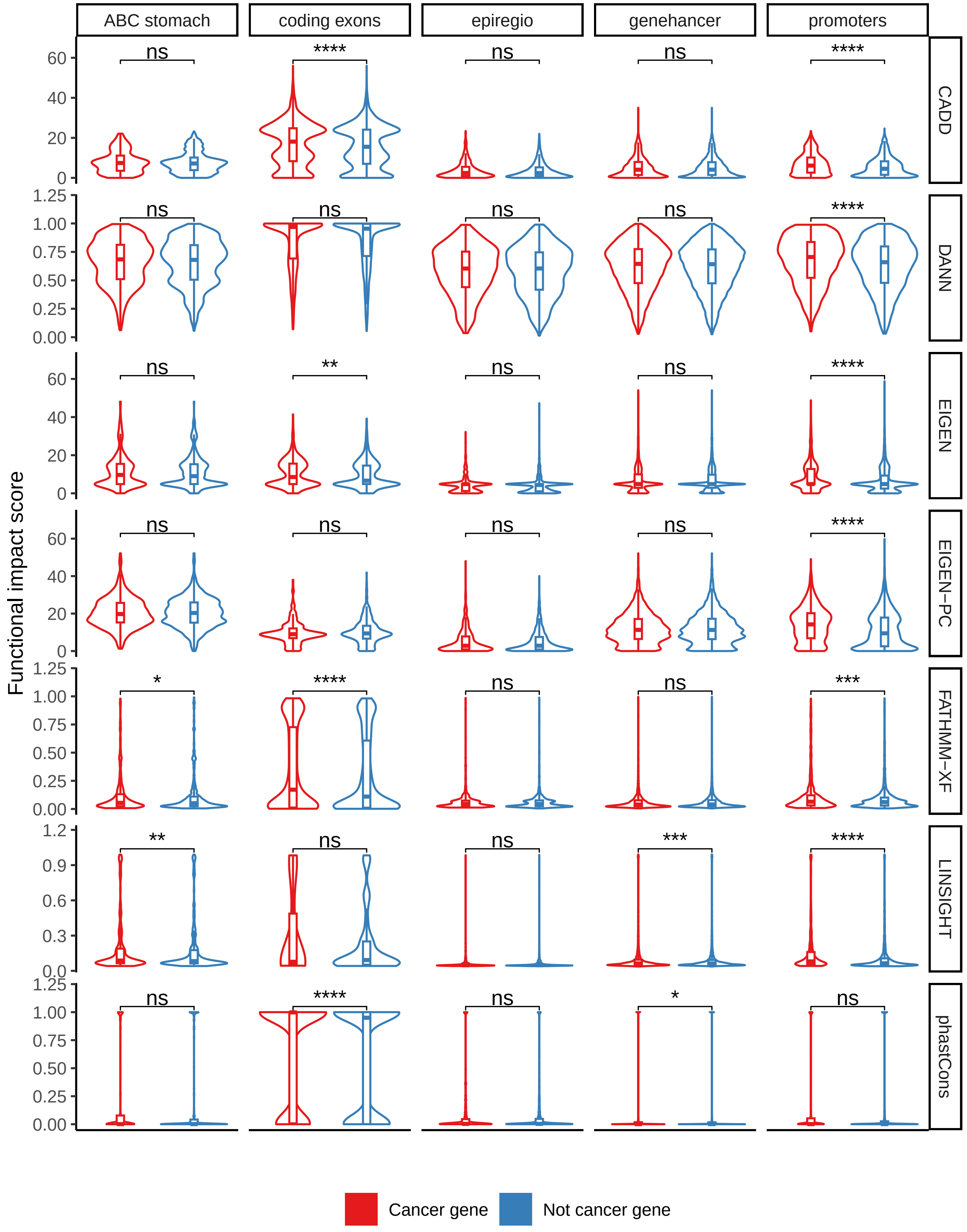


**Supplementary Figure 19: Impact score distributions per region and score type for the observed mutations in the STAD-US cohort.** Distribution of functional impact scores for cancer (red) and non-cancer (blue) genes separated by association and scoring type. Significance labels indicate the p-value for the one-sided Wilcoxon test checking whether the $\log_{10}$(mutation rate) for the cancer genes $>$ $\log_{10}$(mutation rate) for the non-cancer genes. Significance labels indicate: ns - $p > 0.05$; $*: p \leq 0.05$; $**: p \leq 0.01$; $***: p \leq 0.001$; $****: p \leq 0.0001$.

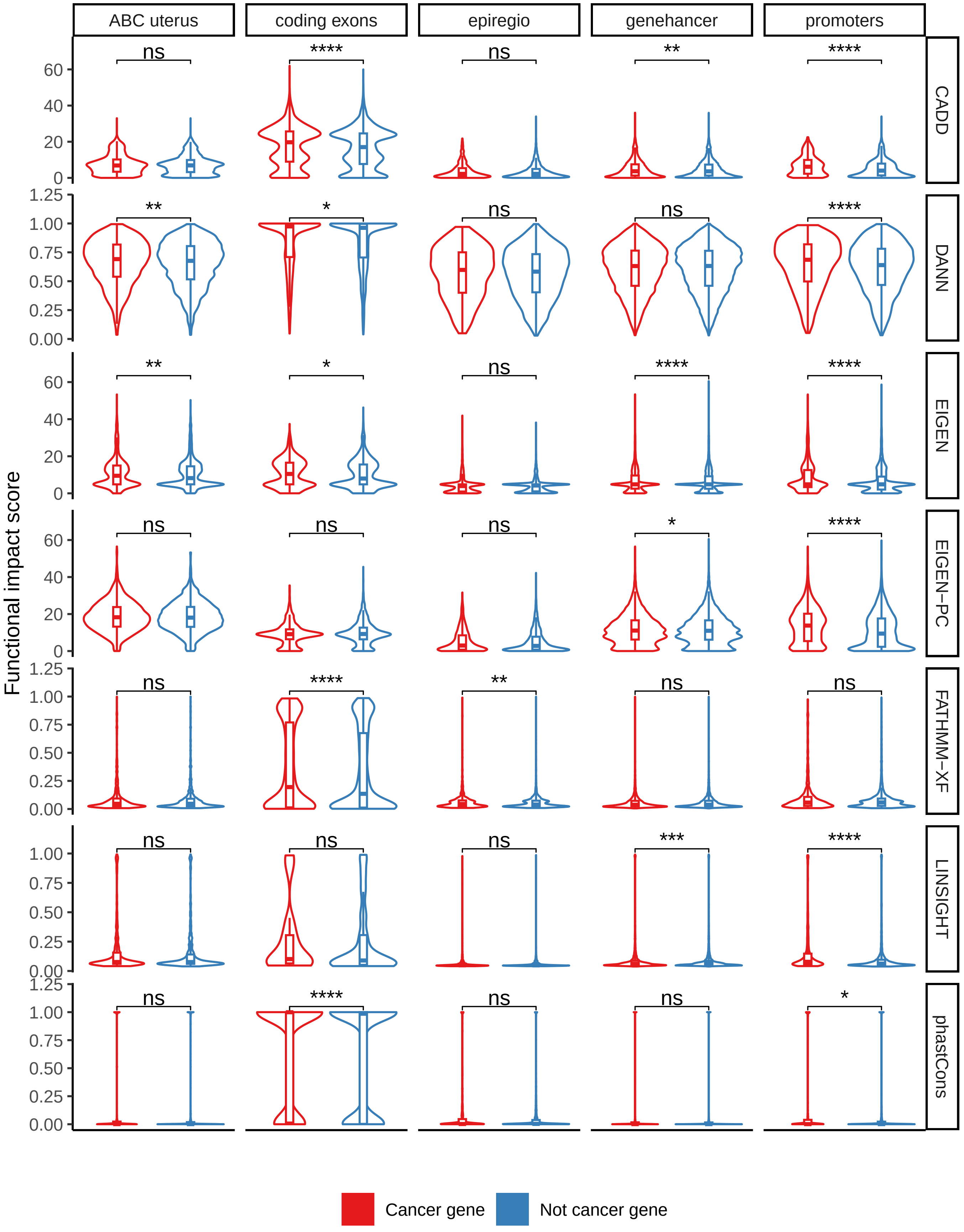


**Supplementary Figure 20: Impact score distributions per region and score type for the observed mutations in the UCEC-US cohort.** Distribution of functional impact scores for cancer (red) and non-cancer (blue) genes separated by association and scoring type. Significance labels indicate the p-value for the one-sided Wilcoxon test checking whether the $\log_{10}$(mutation rate) for the cancer genes $>$ $\log_{10}$(mutation rate) for the non-cancer genes. Significance labels indicate: ns - $p > 0.05$; $* : p \leq 0.05$; $** : p \leq 0.01$; $*** : p \leq 0.001$; $**** : p \leq 0.0001$.

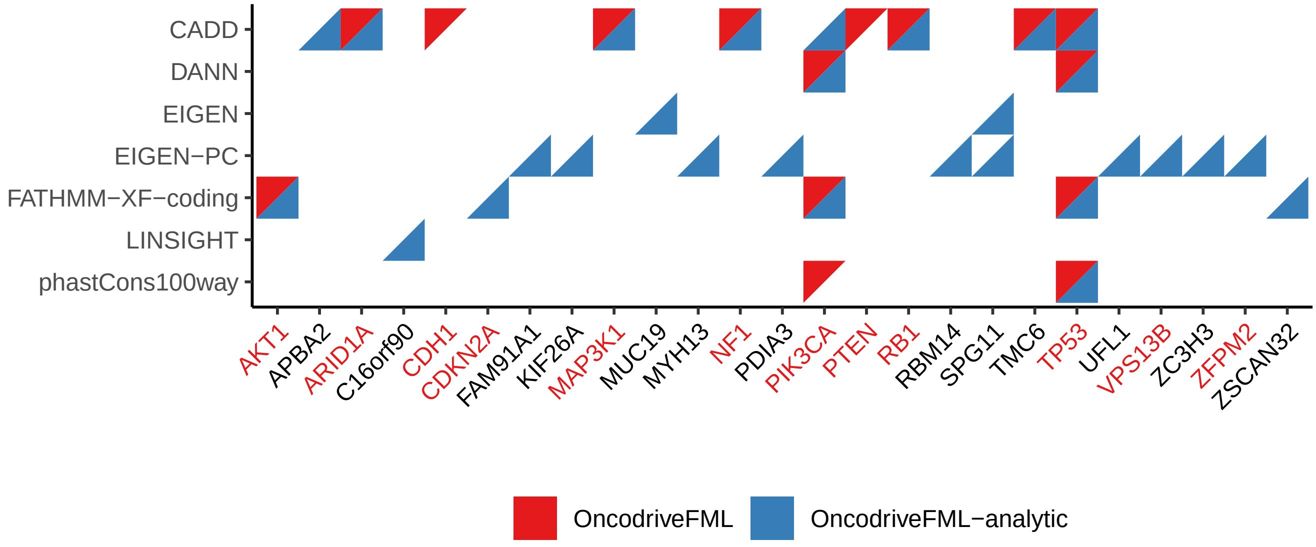


**Supplementary Figure 21: Detected genes across scoring approaches in the analysis of coding exonic sequences with mutations from the BASIS cohort.** Genes (x-axis) showing an enrichment (adjusted-pval $\leq$ 0.05) of high functional impact mutations across different mutation scoring approaches (y-axis) detected by either OncodriveFML (red triangles) or OncodriveFML-analytic (blue triangles). Gene names in red are listed as cancer genes.

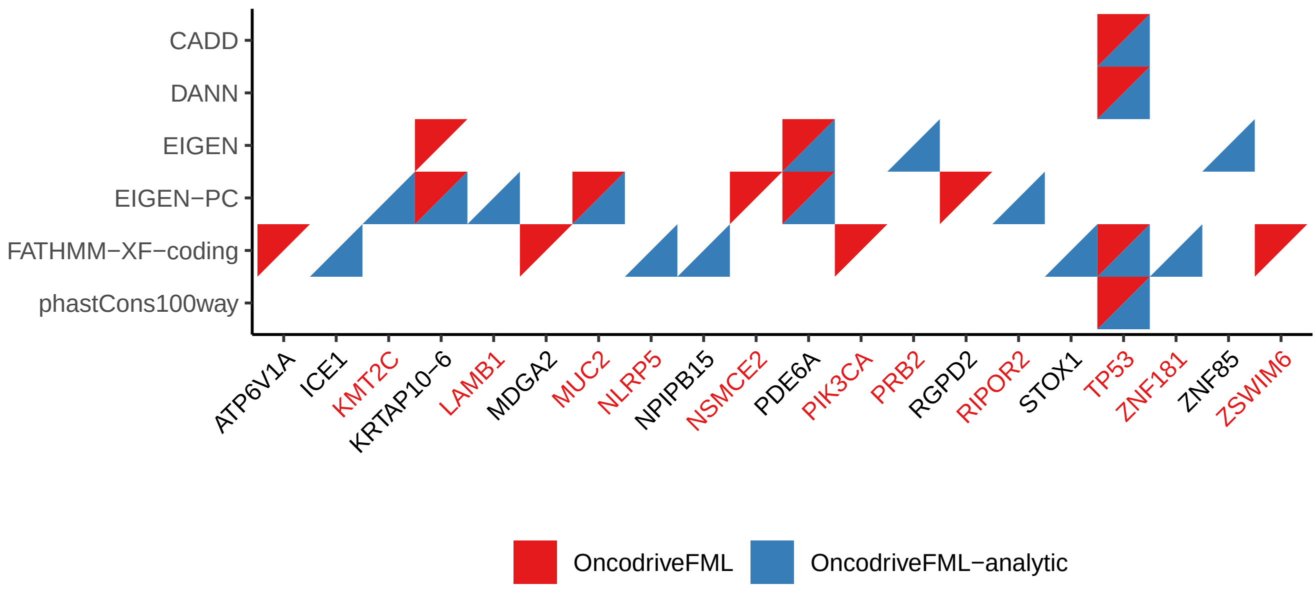


**Supplementary Figure 22: Detected genes across scoring approaches in the analysis of coding exonic sequences with mutations from the BRCA-US cohort.** Genes (x-axis) showing an enrichment (adjusted-pval $\leq$ 0.05) of high functional impact mutations across different mutation scoring approaches (y-axis) detected by either OncodriveFML (red triangles) or OncodriveFML-analytic (blue triangles). Gene names in red are listed as cancer genes.

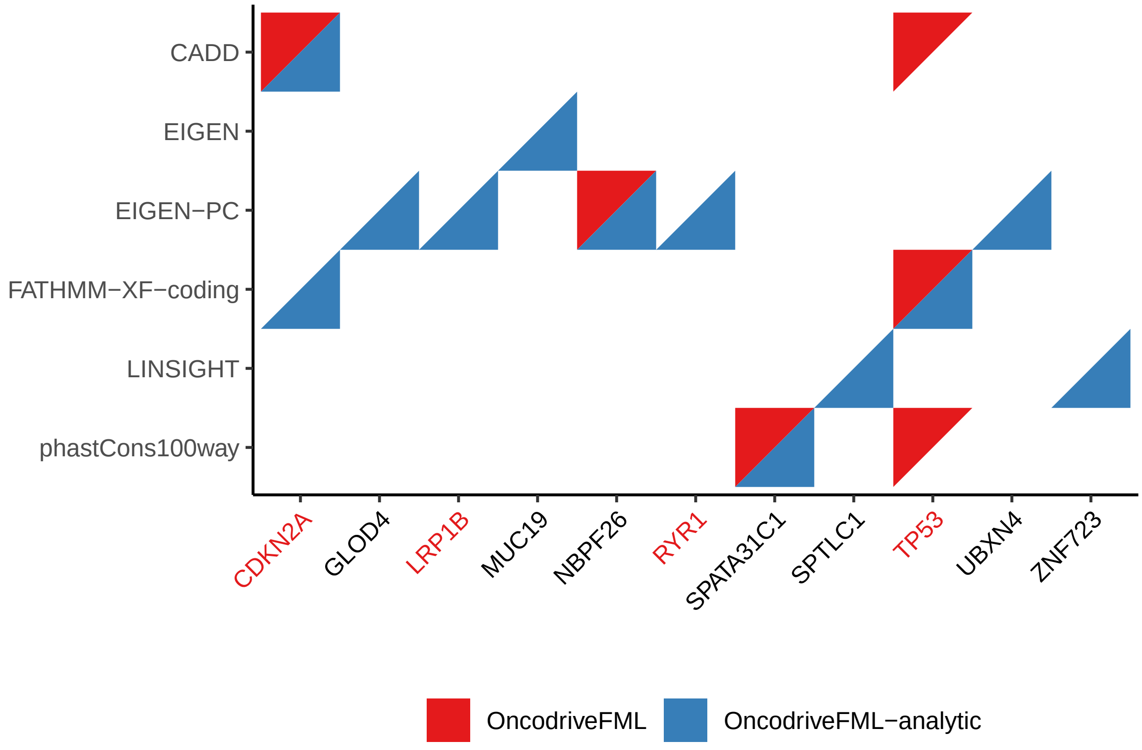


**Supplementary Figure 23: Detected genes across scoring approaches in the analysis of coding exonic sequences with mutations from the HNSC-US cohort.** Genes (x-axis) showing an enrichment (adjusted-pval $\leq$ 0.05) of high functional impact mutations across different mutation scoring approaches (y-axis) detected by either OncodriveFML (red triangles) or OncodriveFML-analytic (blue triangles). Gene names in red are listed as cancer genes.

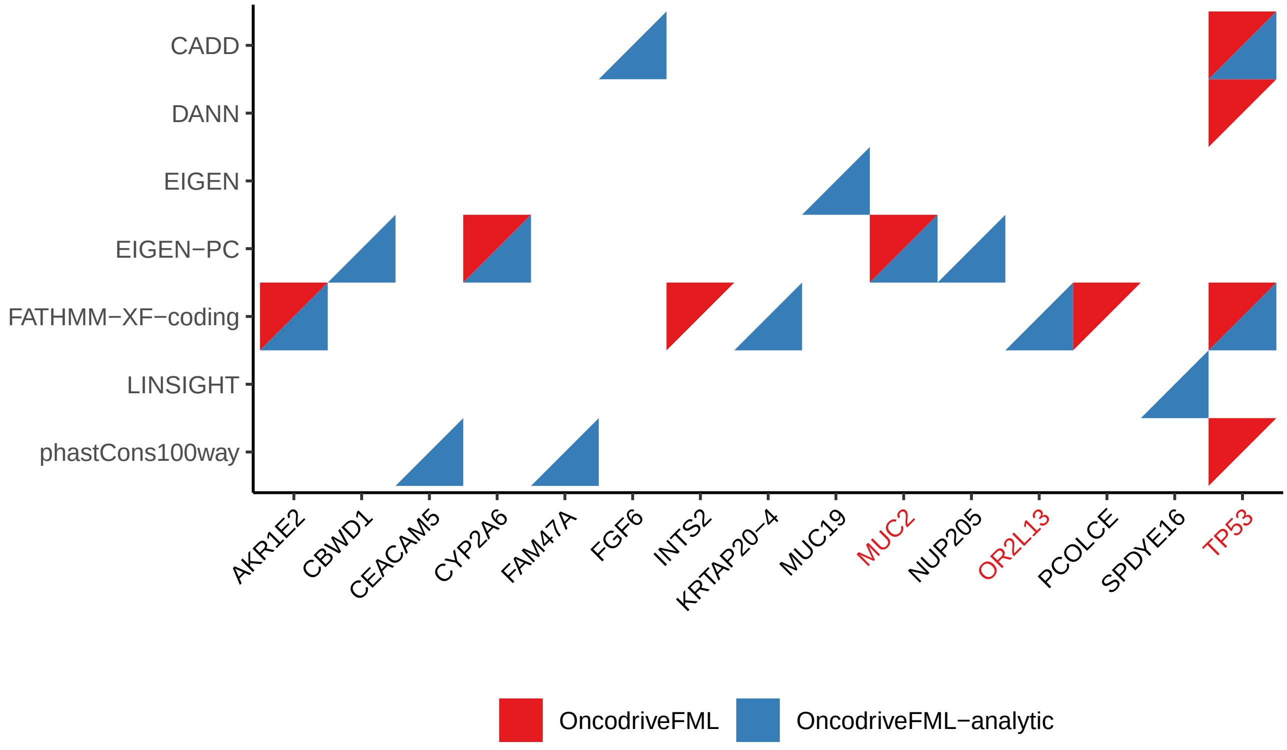


**Supplementary Figure 24: Detected genes across scoring approaches in the analysis of coding exonic sequences with mutations from the LIHC-US cohort.** Genes (x-axis) showing an enrichment (adjusted-pval $\leq$ 0.05) of high functional impact mutations across different mutation scoring approaches (y-axis) detected by either OncodriveFML (red triangles) or OncodriveFML-analytic (blue triangles). Gene names in red are listed as cancer genes.

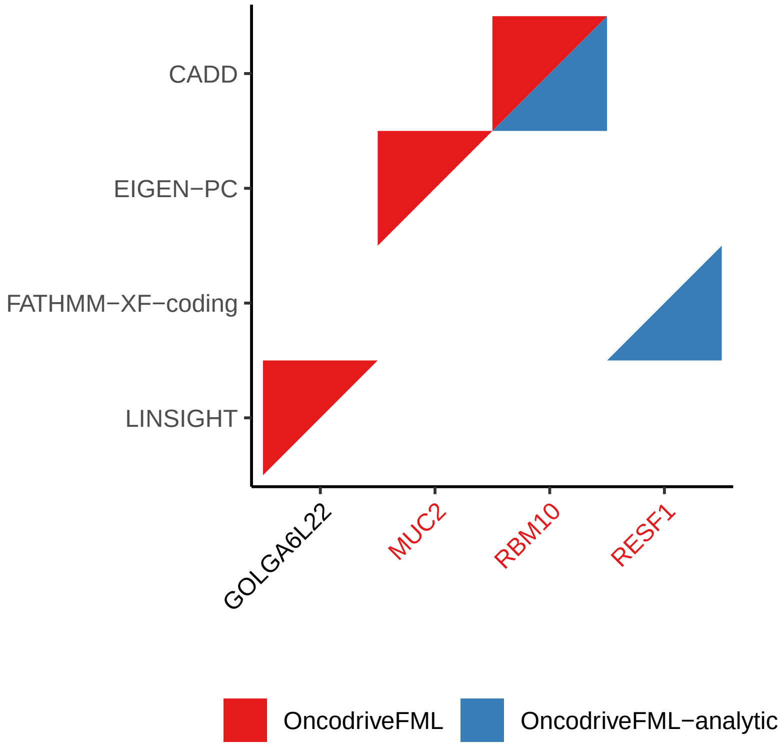


**Supplementary Figure 25: Detected genes across scoring approaches in the analysis of coding exonic sequences with mutations from the LUAD-US cohort.** Genes (x-axis) showing an enrichment (adjusted-pval $\leq$ 0.05) of high functional impact mutations across different mutation scoring approaches (y-axis) detected by either OncodriveFML (red triangles) or OncodriveFML-analytic (blue triangles). Gene names in red are listed as cancer genes.

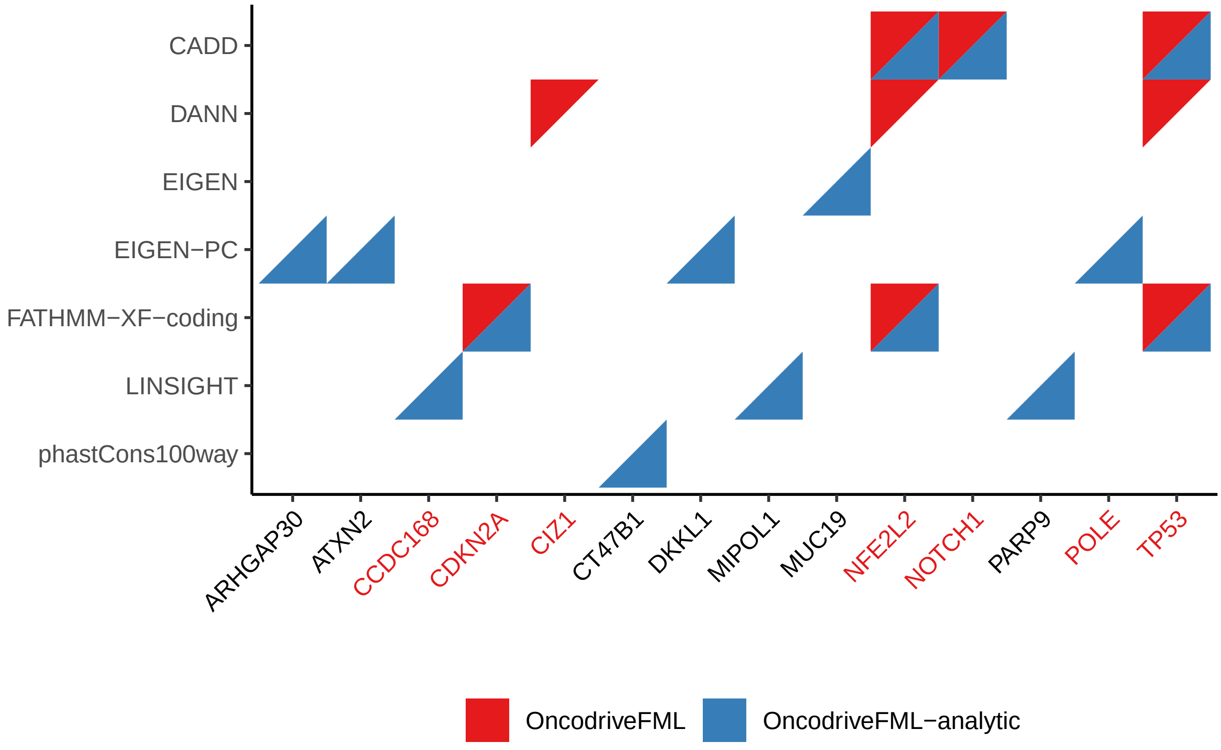


**Supplementary Figure 26: Detected genes across scoring approaches in the analysis of coding exonic sequences with mutations from the LUSC-US cohort.** Genes (x-axis) showing an enrichment (adjusted-pval $\leq$ 0.05) of high functional impact mutations across different mutation scoring approaches (y-axis) detected by either OncodriveFML (red triangles) or OncodriveFML-analytic (blue triangles). Gene names in red are listed as cancer genes.

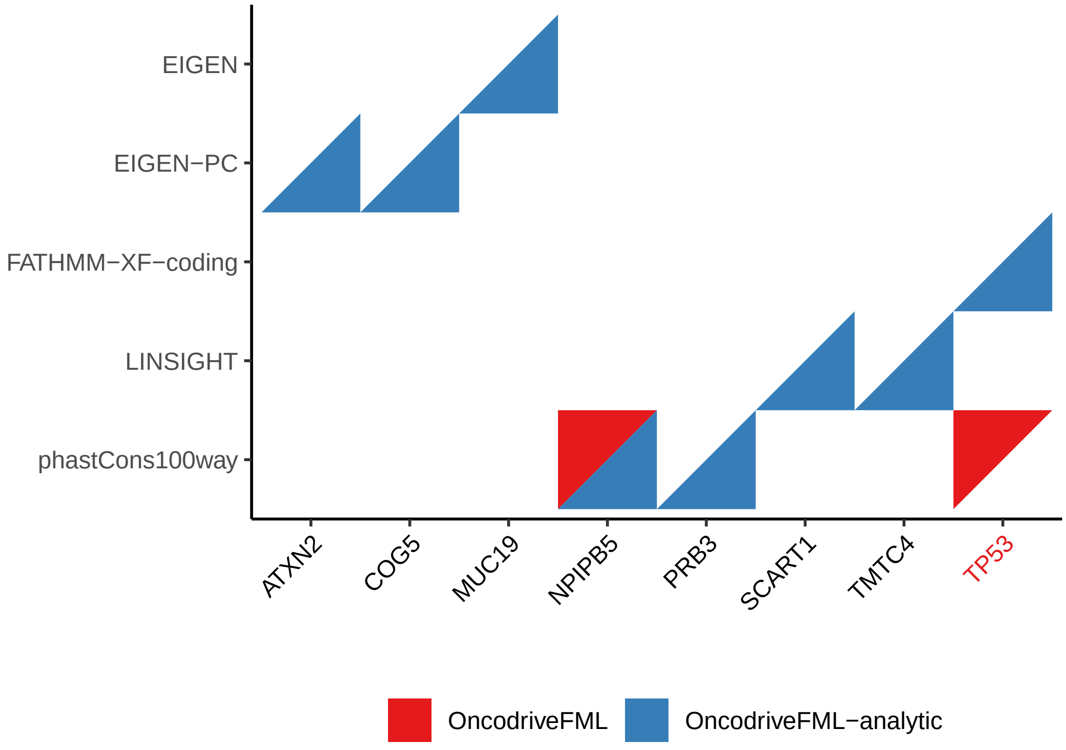


**Supplementary Figure 27: Detected genes across scoring approaches in the analysis of coding exonic sequences with mutations from the STAD-US cohort.** Genes (x-axis) showing an enrichment (adjusted-pval $\leq$ 0.05) of high functional impact mutations across different mutation scoring approaches (y-axis) detected by either OncodriveFML (red triangles) or OncodriveFML-analytic (blue triangles). Gene names in red are listed as cancer genes.

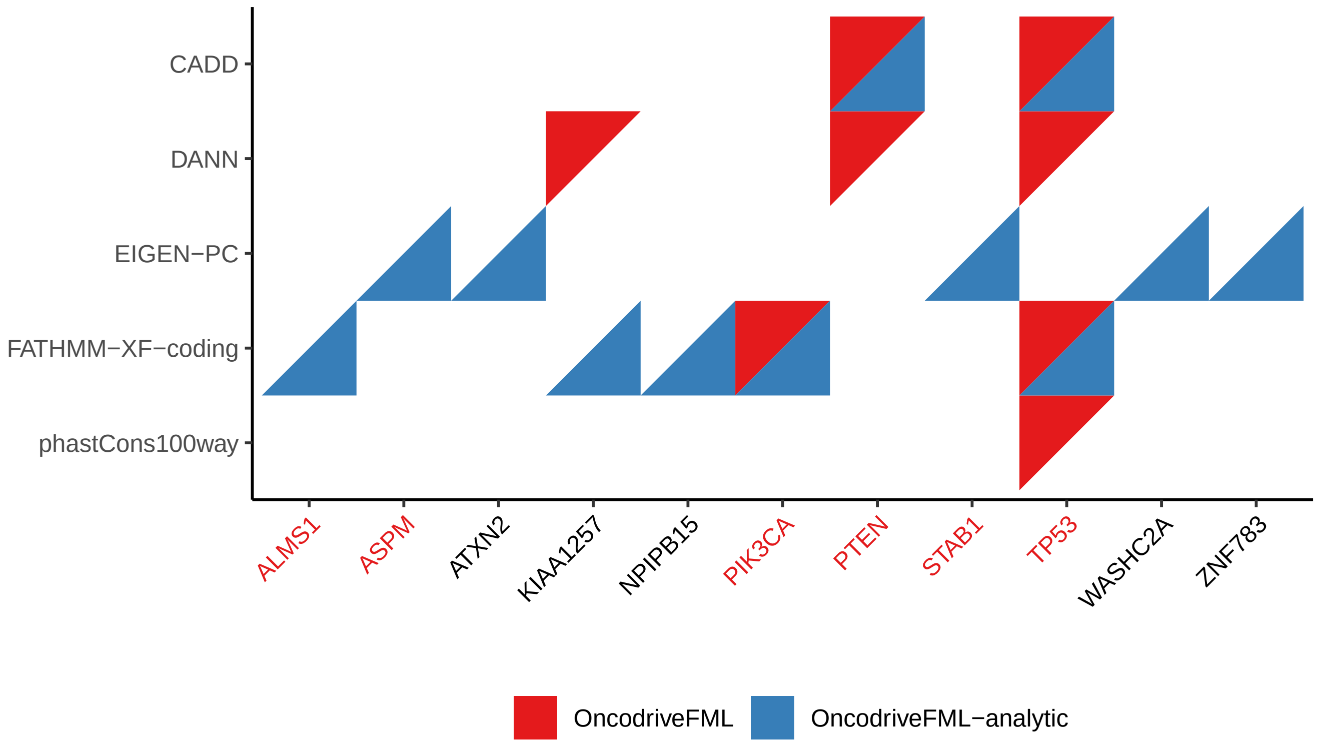


**Supplementary Figure 28: Detected genes across scoring approaches in the analysis of coding exonic sequences with mutations from the UCEC-US cohort.** Genes (x-axis) showing an enrichment (adjusted-pval $\leq$ 0.05) of high functional impact mutations across different mutation scoring approaches (y-axis) detected by either OncodriveFML (red triangles) or OncodriveFML-analytic (blue triangles). Gene names in red are listed as cancer genes.

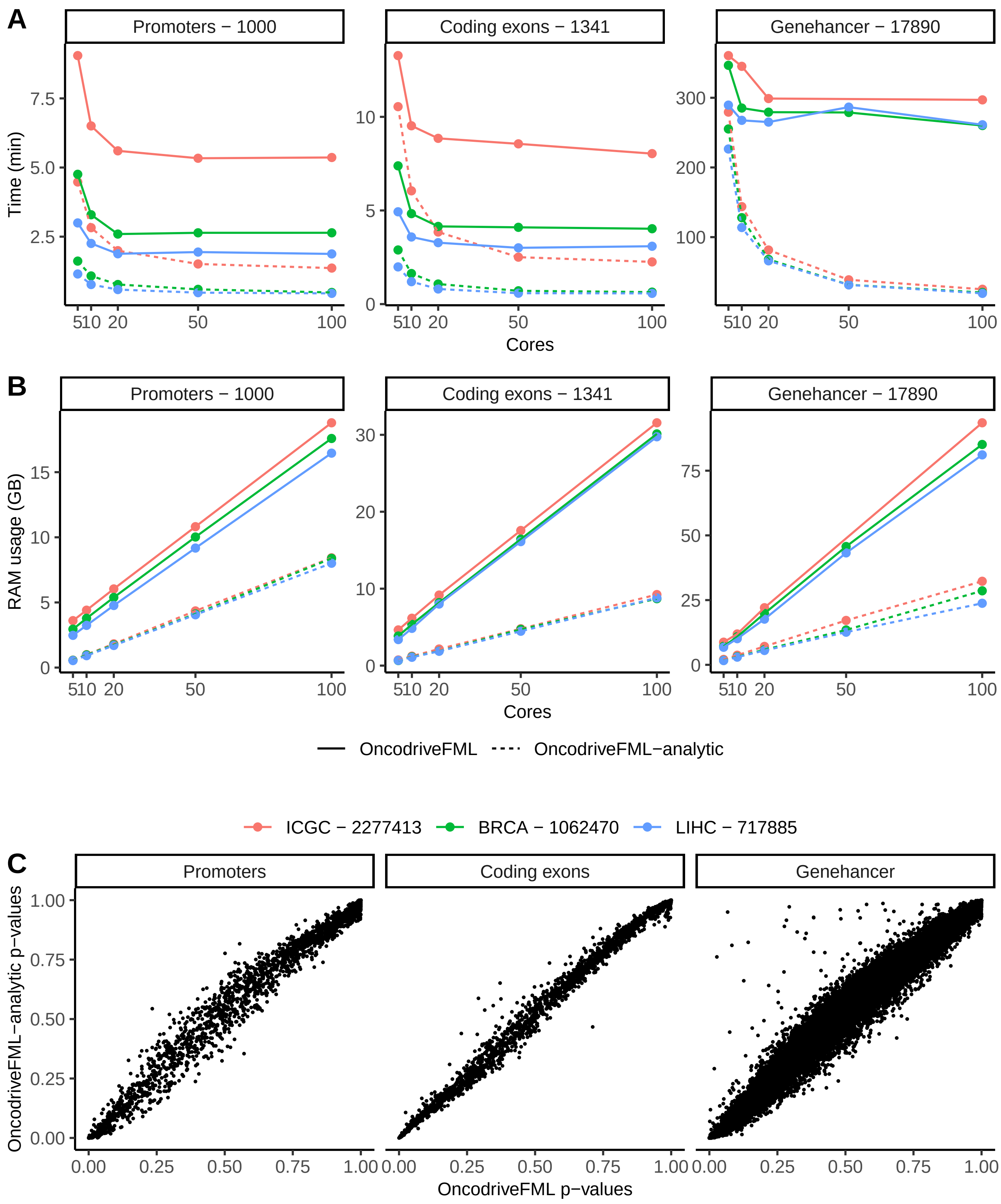


**Supplementary Figure 29: Computational benchmark of OncodriveFML-analytic and OncodriveFML.** Elapsed time in minutes **(A)** and RAM usage in GB **(B)** (y-axis) versus numbers of cores (x-axis) used by OncodriveFML-analytic (solid line) or OncodriveFML (dahsed line) in regions of increasing sizes. Region median sizes are indicated in the facet titles. Colors indicate the used cohorts in the benchmark together with the total number of observed mutations in each cohort. **(C)** Comparison of each gene's p-values between OncodriveFML (x-axis) and OncodriveFML-analytic (y-axis).

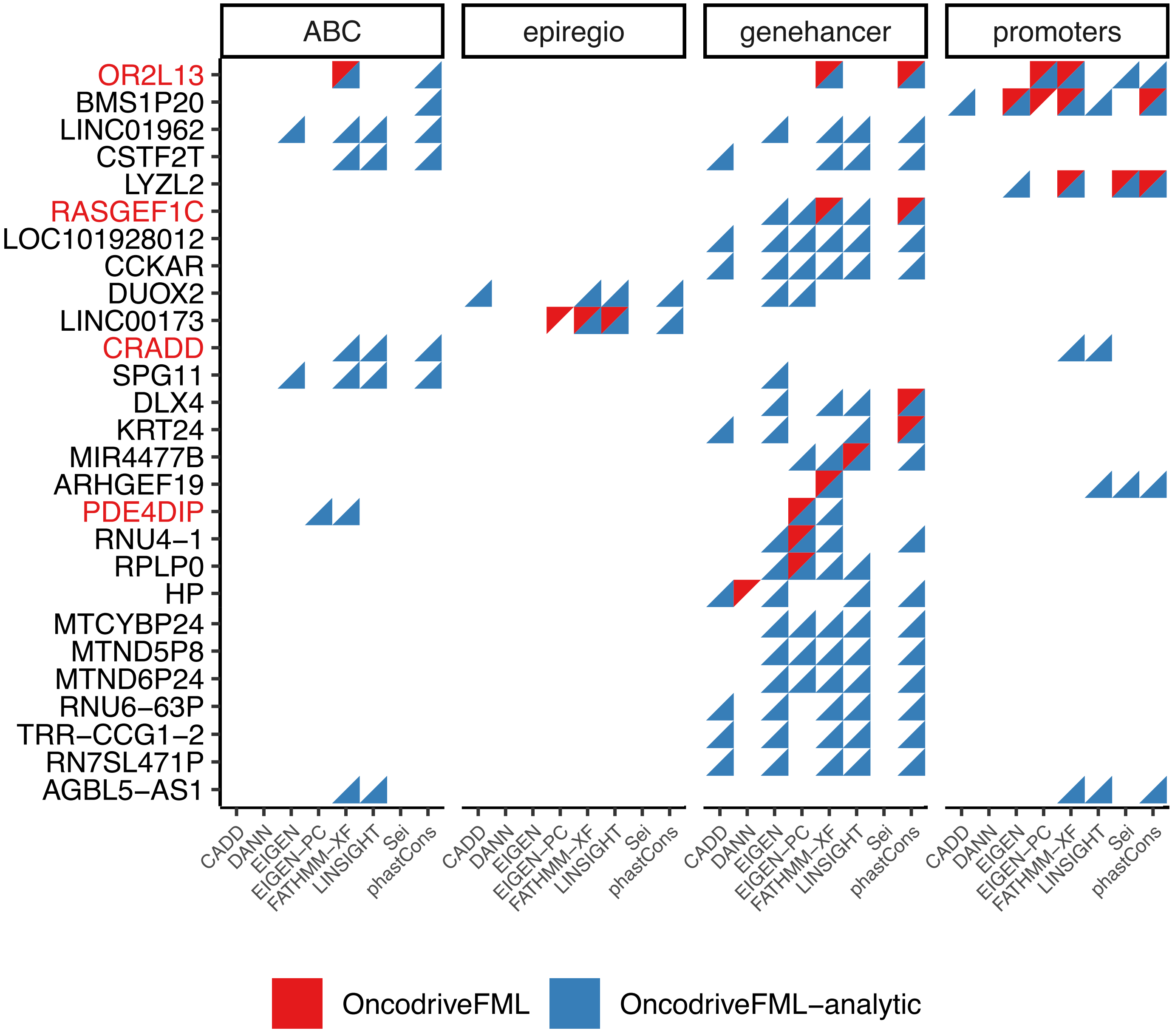


**Supplementary Figure 30: Detected genes across CRR-score combinations in the TCGA BRCA cancer cohort.** Genes (y-axis) showing an enrichment (adjusted-pval $\leq 0.05$) of FI CRVs in different score (x-axis) and association (facet) combinations detected by either OncodriveFML (red triangles) or OncodriveFML-analytic (blue triangles). Only genes detected in at least five or more CRR-score combinations and/or enrichment method are shown. Gene names in red are listed as cancer genes.

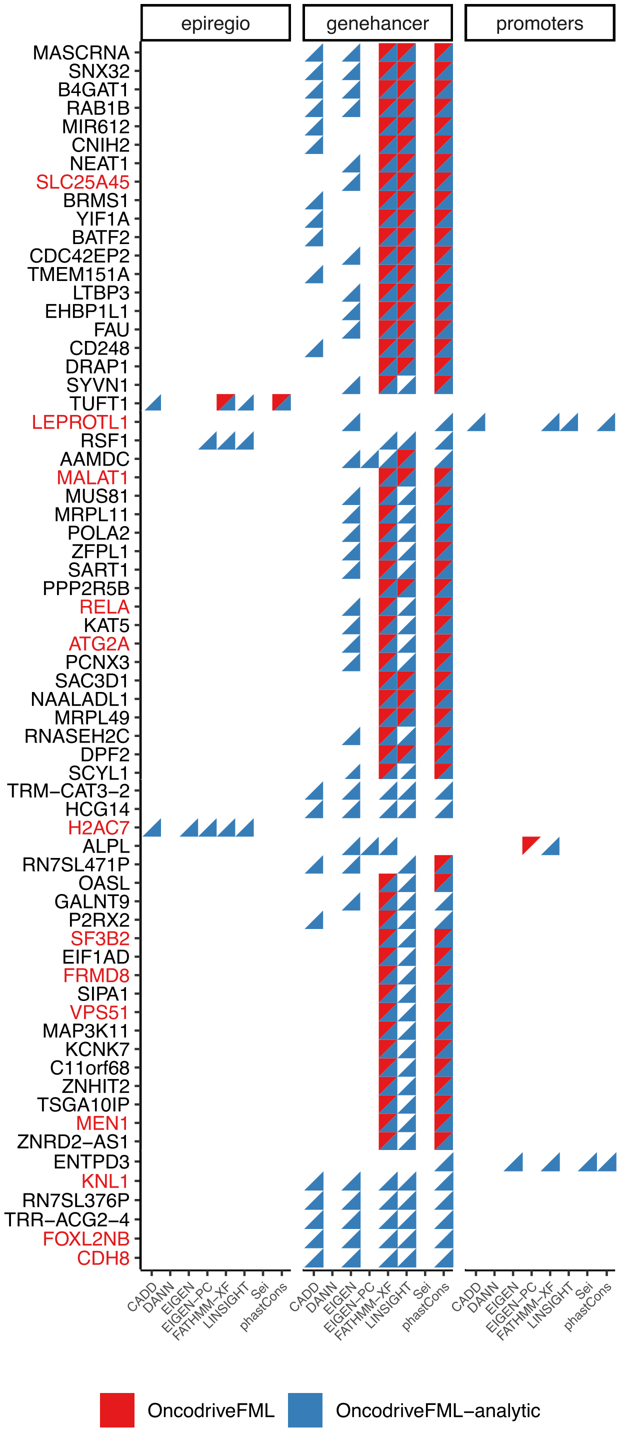


**Supplementary Figure 31: Detected genes across CRR-score combinations in the TCGA HNSC cancer cohort.** Genes (y-axis) showing an enrichment (adjusted-pval $\leq 0.05$) of FI CRVs in different score (x-axis) and association (facet) combinations detected by either OncodriveFML (red triangles) or OncodriveFML-analytic (blue triangles). Only genes detected in at least five or more CRR-score combinations and/or enrichment method are shown. Gene names in red are listed as cancer genes.

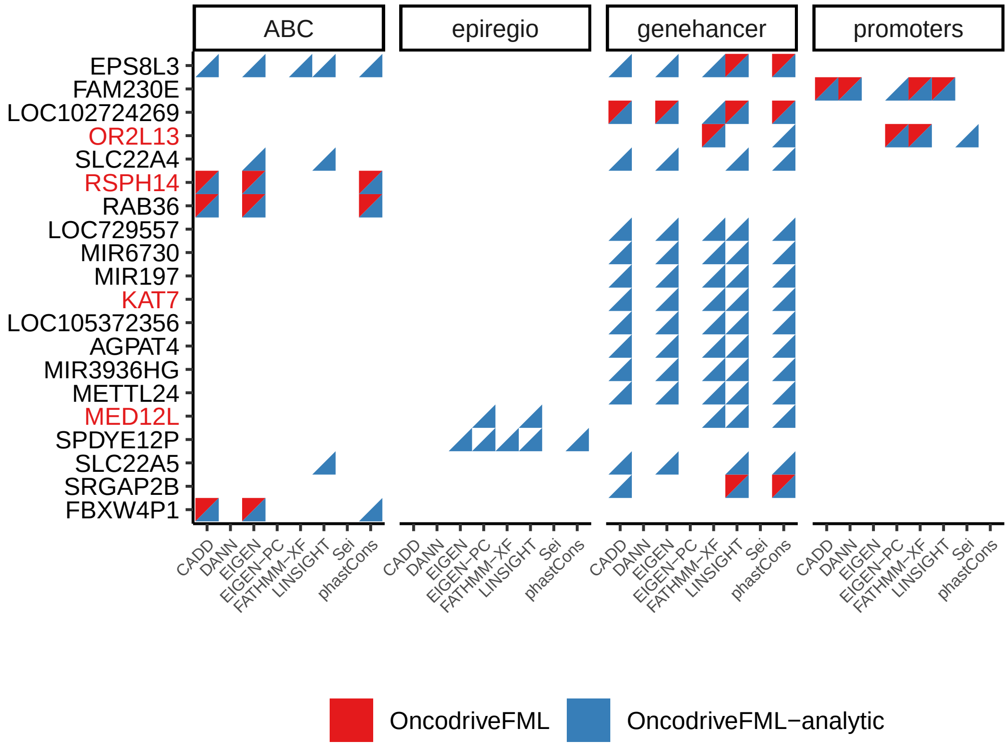


**Supplementary Figure 32: Detected genes across CRR-score combinations in the TCGA LIHC cancer cohort.** Genes (y-axis) showing an enrichment (adjusted-pval $\leq 0.05$) of FI CRVs in different score (x-axis) and association (facet) combinations detected by either OncodriveFML (red triangles) or OncodriveFML-analytic (blue triangles). Only genes detected in at least five or more CRR-score combinations and/or enrichment method are shown. Gene names in red are listed as cancer genes.

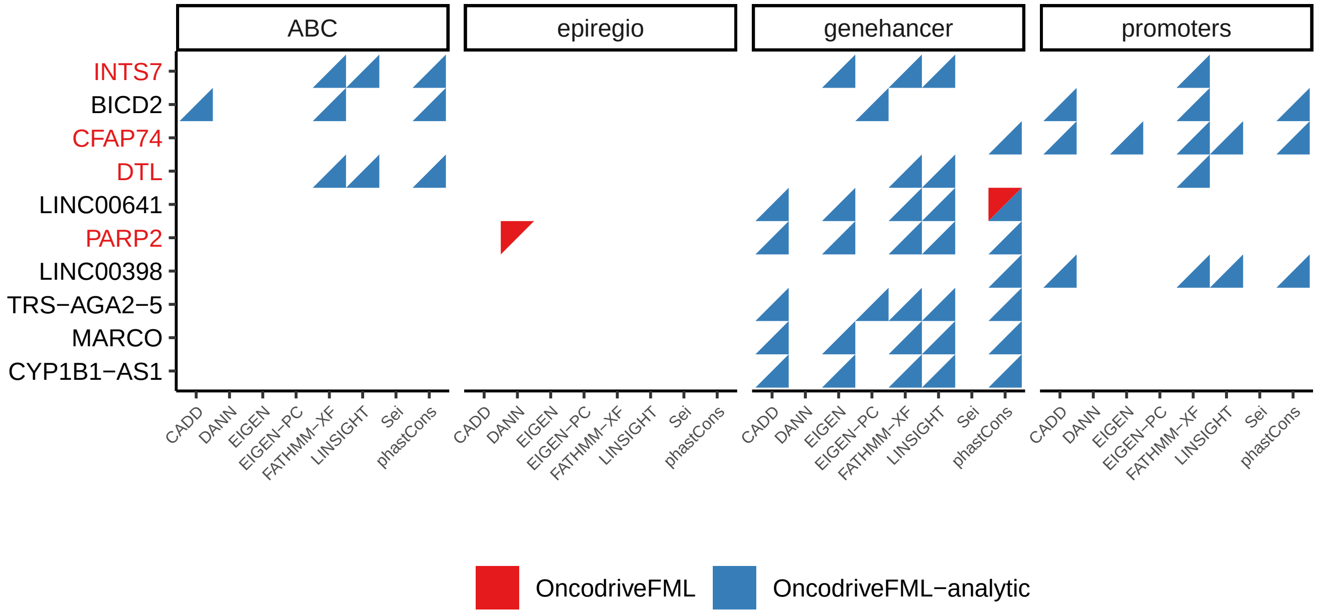


**Supplementary Figure 33: Detected genes across CRR-score combinations in the TCGA LUAD cancer cohort.** Genes (y-axis) showing an enrichment (adjusted-pval $\leq 0.05$) of FI CRVs in different score (x-axis) and association (facet) combinations detected by either OncodriveFML (red triangles) or OncodriveFML-analytic (blue triangles). Only genes detected in at least five or more CRR-score combinations and/or enrichment method are shown. Gene names in red are listed as cancer genes.

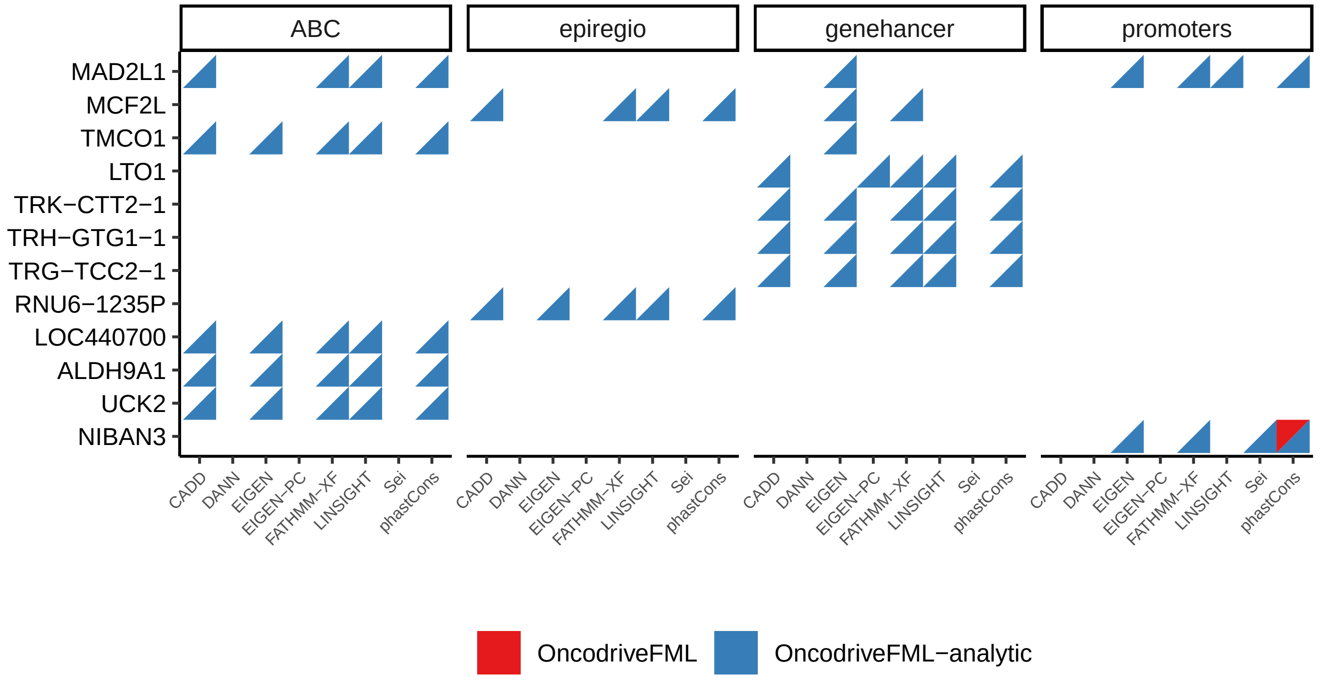


**Supplementary Figure 34: Detected genes across CRR-score combinations in the TCGA LUSC cancer cohort.** Genes (y-axis) showing an enrichment (adjusted-pval $\leq$ 0.05) of FI CRVs in different score (x-axis) and association (facet) combinations detected by either OncodriveFML (red triangles) or OncodriveFML-analytic (blue triangles). Only genes detected in at least five or more CRR-score combinations and/or enrichment method are shown. Gene names in red are listed as cancer genes.

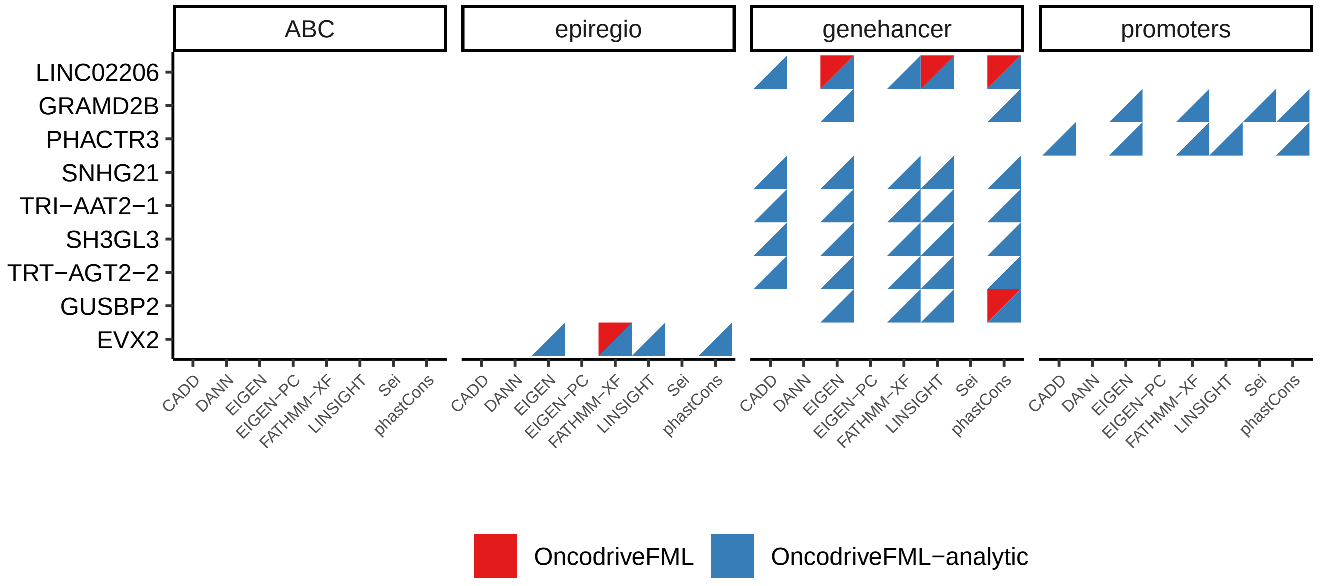


**Supplementary Figure 35: Detected genes across CRR-score combinations in the TCGA STAD cancer cohort.** Genes (y-axis) showing an enrichment (adjusted-pval $\leq 0.05$) of FI CRVs in different score (x-axis) and association (facet) combinations detected by either OncodriveFML (red triangles) or OncodriveFML-analytic (blue triangles). Only genes detected in at least five or more CRR-score combinations and/or enrichment method are shown. Gene names in red are listed as cancer genes.

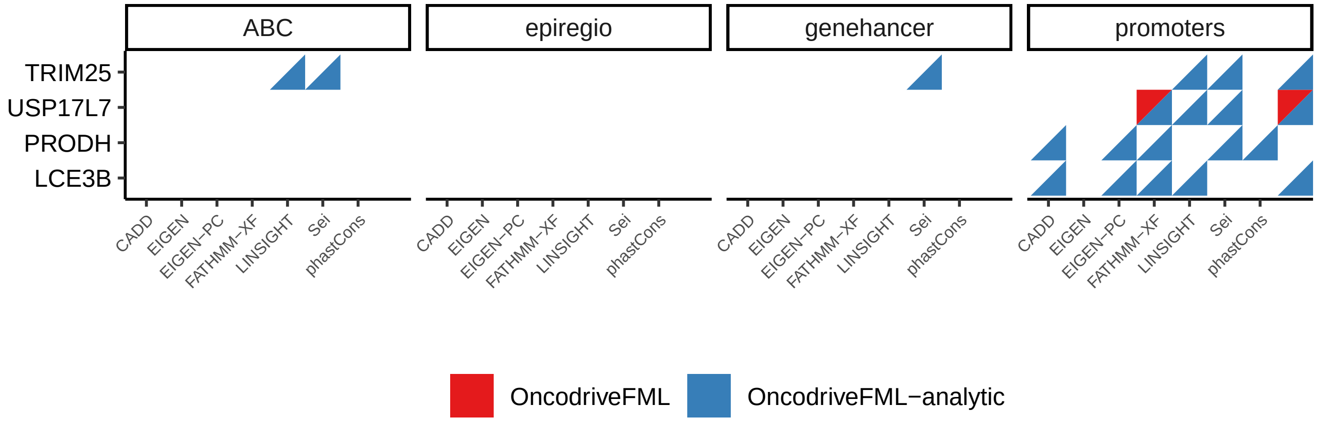


**Supplementary Figure 36: Detected genes across CRR-score combinations in the TCGA UCEC cancer cohort.** Genes (y-axis) showing an enrichment (adjusted-pval $\leq 0.05$) of FI CRVs in different score (x-axis) and association (facet) combinations detected by either OncodriveFML (red triangles) or OncodriveFML-analytic (blue triangles). Only genes detected in at least five or more CRR-score combinations and/or enrichment method are shown. Gene names in red are listed as cancer genes.

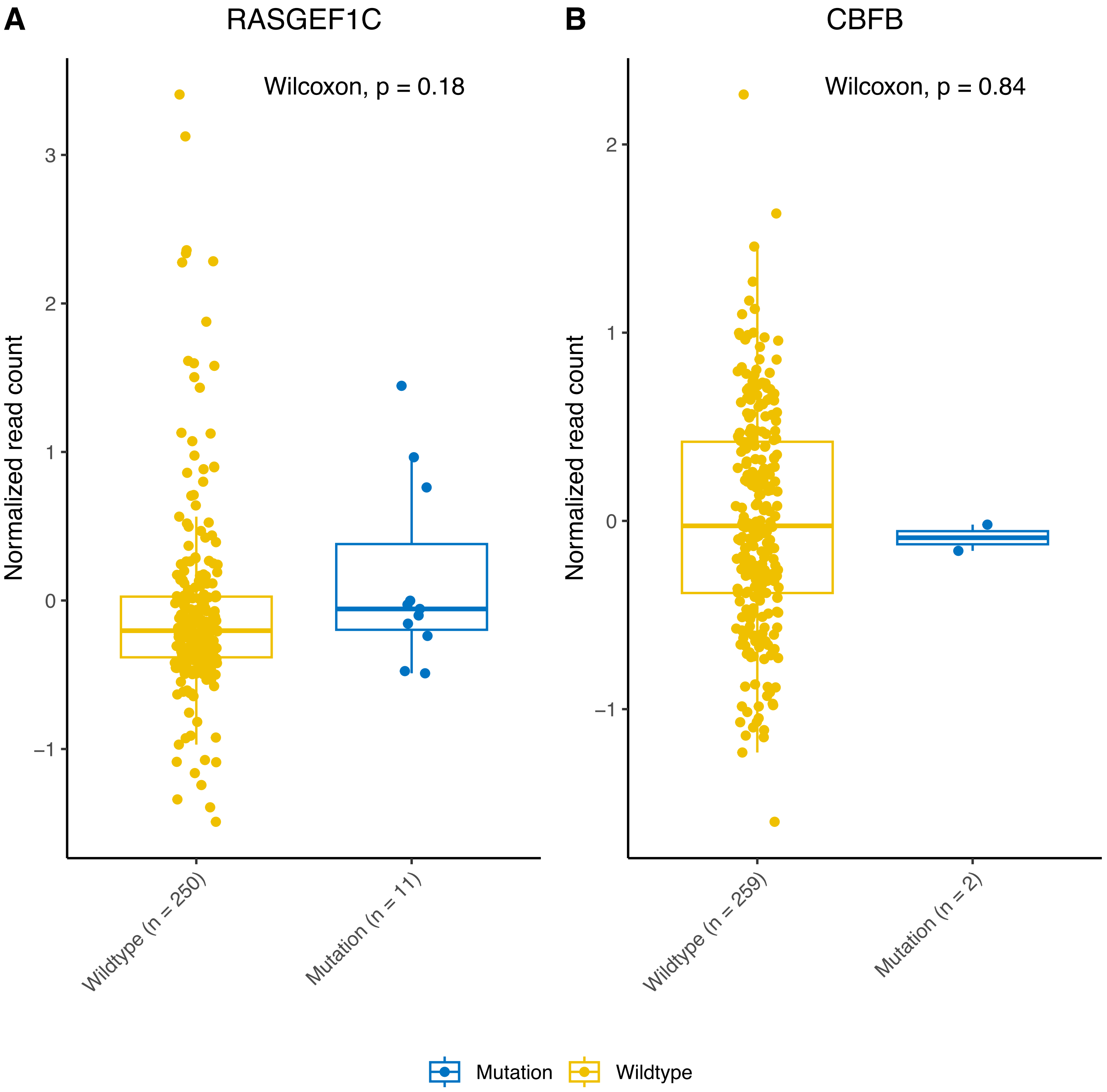


**Supplementary Figure 37: CBFB and RASGEF1C do not show changes in their expression.** Boxplots comparing the normalized expression of RASGEF1C (left) and CBFB (right) stratified by patients showing mutations in their CRRs.

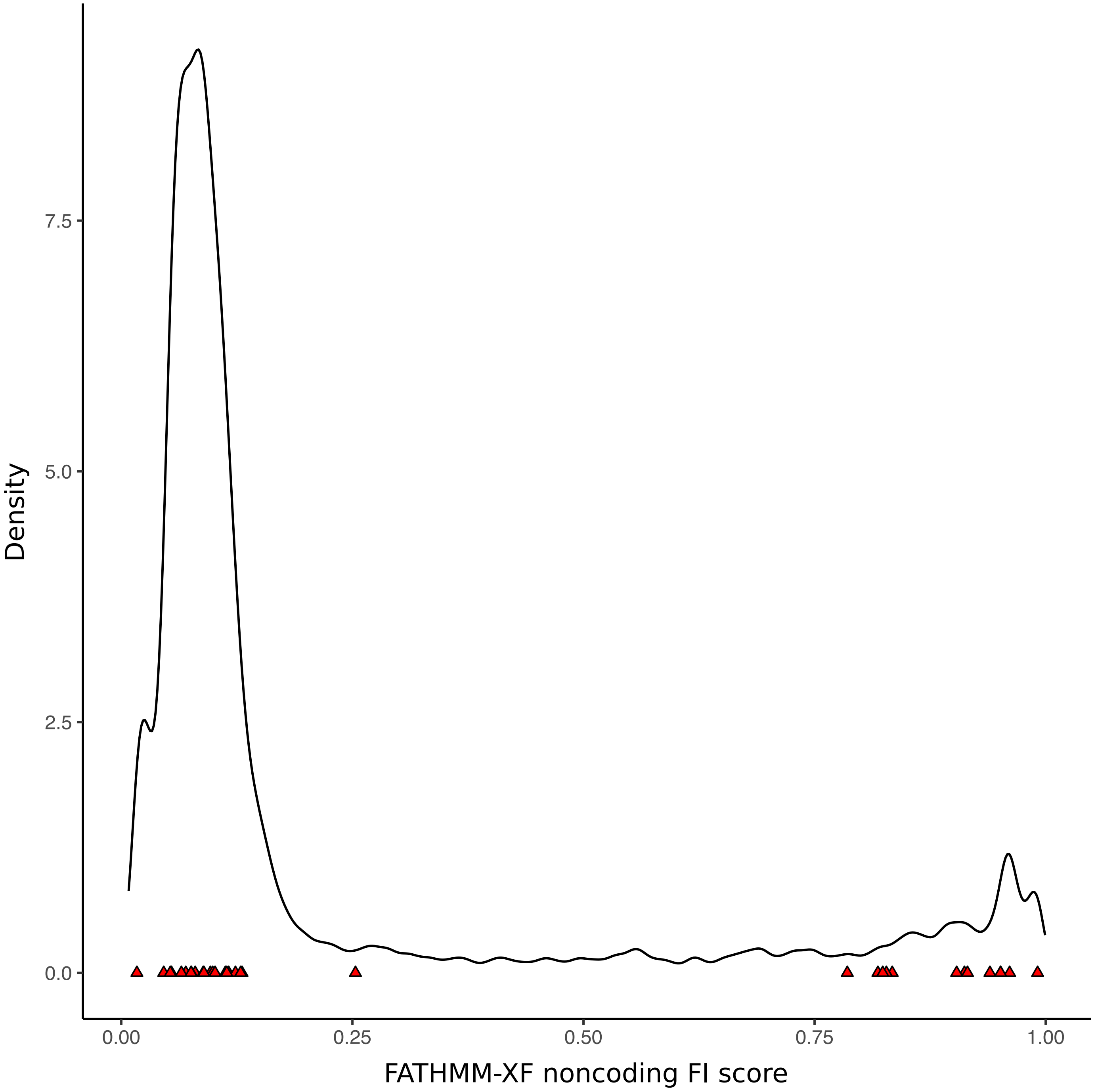


**Supplementary Figure 38: Distribution of functional impact scores in the ABC-derived BRCA1 CRRs.** Density plot showing the distribution of possible functional impact scores within the ABC-derived BRCA1 CRRs. Red triangles show the scores of the observed SNPs in the CRRs of BRCA1.

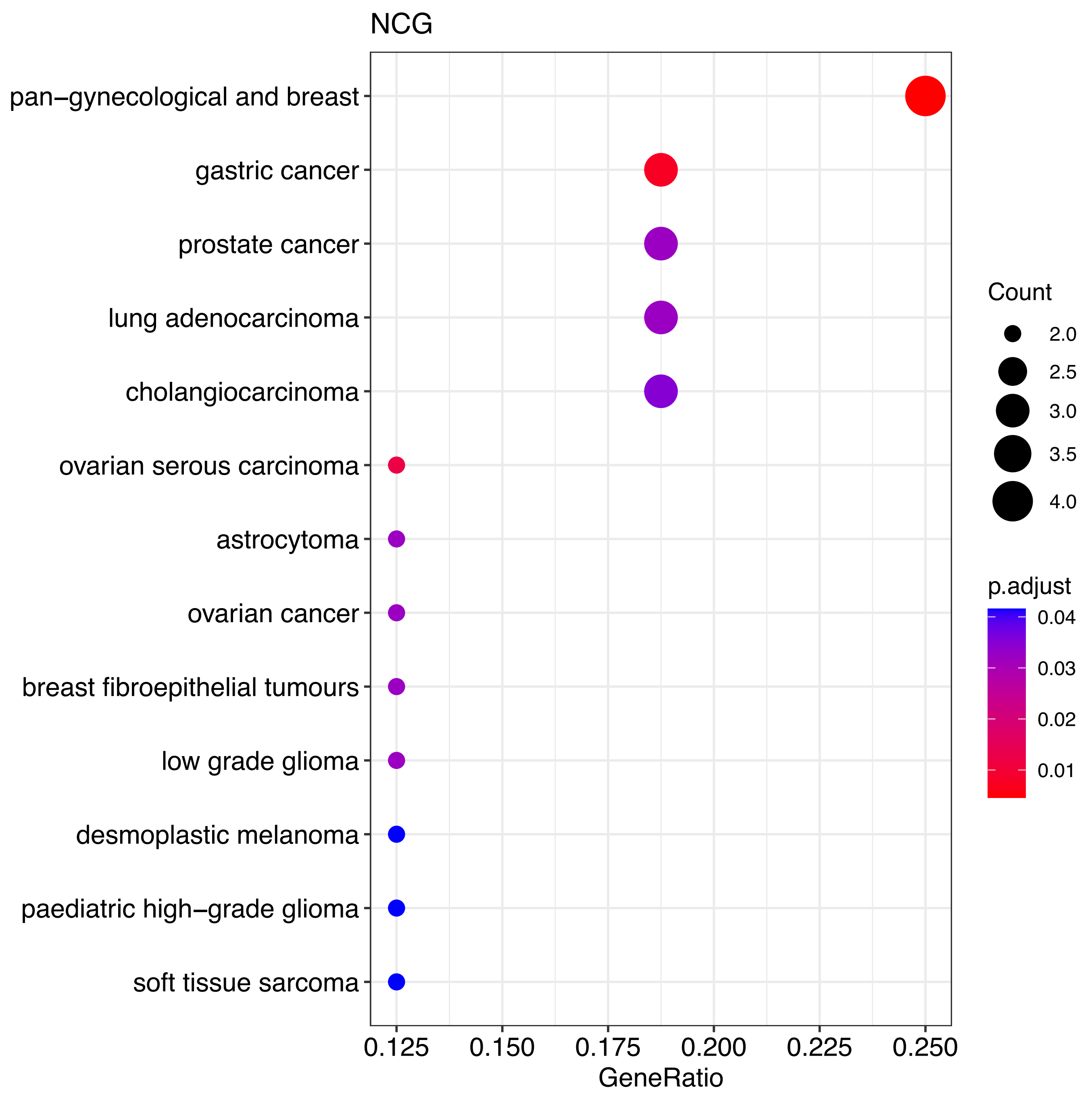


**Supplementary Figure 39: Enrichment of breast-cancer-related genes showing an enrichment of FI SNPs in their CRRs.** Gene set enrichment analysis using the Network of Cancer Genes ontology on the genes with an enrichment of FI SNPs in their CRRs.

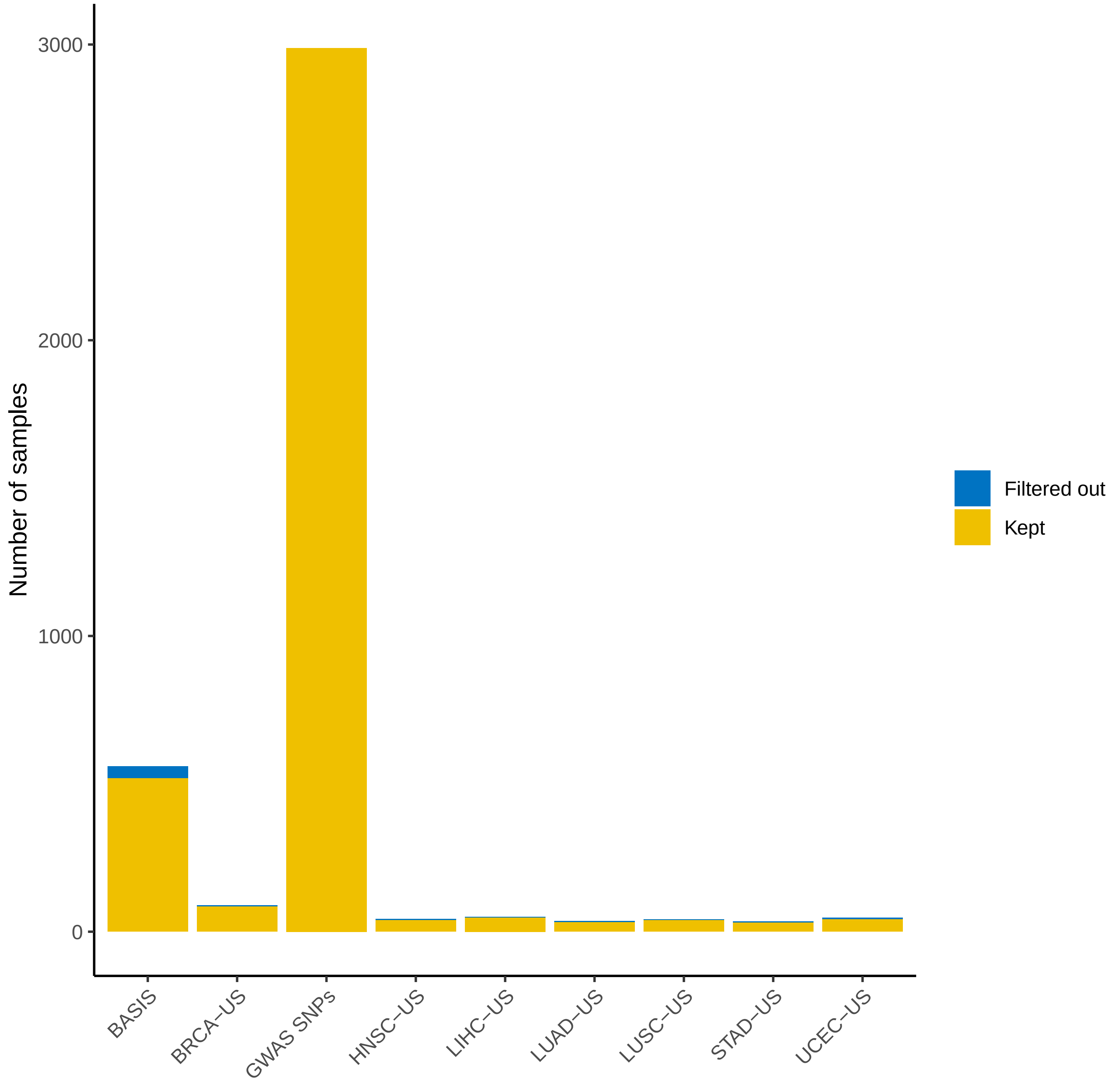


**Supplementary Figure 40: Numbers of samples per cohort before and after filtering.** Number of samples (y-axis) by cohort (x-axis). Samples that passed the filtering step are colored in yellow, while filtered out samples are colored in blue.
